# An environment to genome control loop using RNA interference processing of secreted tRNAs may regulates the *C. elegans* chemo-sensory behavior

**DOI:** 10.1101/2022.06.22.496966

**Authors:** Jean-Jacques Remy

## Abstract

Alanine tRNAs (UGC) control the development of the innate and the environment-modulated acquired *C. elegans* chemo-attractive responses. Some Ala-tRNA isomers are required for the development of the chemo-attractive behavior (dev-tRNAs), while others (odor-tRNAs) are made as life-term olfactory imprints of early larval odor-exposures.

dev-tRNAs and odor-tRNAs biosynthesis respectively require the tRNA modifying Elongator complex sub-units ELPC-3 and ELPC-1: while *elpc-3* mutants are chemo-attraction deficients, *elpc-1* mutants do not synthesize odor-tRNAs imprints.

Feeding wild-type dev-tRNAs restore a wild-type behavior in *elpc-3* mutants. Feeding purified odor-tRNAs enhances odor responses (positive imprinting) in adult wild-type worms, while it decreases odor responses (negative imprinting) in adult imprinting deficient *elpc-1* mutants.

Both positive and negative imprinting can be stably inherited in worm populations. Crossing experiments indicate that both behavioral phenotypes segregate as monogenic monoallelic alterations, following Mendelian inheritance rules.

Co-culture and food conditioning suggest the developmental and the odor-specific regulatory Ala-tRNAs are released in worms environment. Commensal naive acquire odor-specific imprinting from odor-experienced, while co-culture together with wild-type animals fully rescues the chemo-attractive defects of the *elpc-3* mutants.

Worm to worm communication of imprinting require a number of RNA interference (RNAi) genes as the intestinal RNA transporter SID-2, the initial exogenous RNAi Dicer/RDE-1/DRH-1-2/RDE-4 complexe, and the RNA-dependent RNA polymerase RRF-3. Moreover, a male contribution of the 3’-exonuclease ERI-1 activity determines whether olfactory imprints will be erased or stably fixed and inherited in worms progeny.

The RNAi processing of externalized chemosensory regulatory Ala-tRNAs would generate small interfering tRNAs (si-tRNAs) able to target only tRNA complementary sequences present on worm genome, that is the tDNA genes and the transcription independent extra-TFIIIC sites.

A model of control loop in which olfactory receptor expression levels in chemosensory neurons could be non-genetically but stably regulated via RNAi processing of secreted constitutive or environment-modified Ala-tRNAs is discussed.

## INTRODUCTION

There is ample evidence that epigenetic mechanisms mediate the acquisition and the trans-generational transmission of environment-induced phenotypic changes. Weither inheritability of environmentally induced acquired characteristics play a central role in evolution is still a matter of debate (***West-Eberhard, 2005; Heard and Martienssen, 2014; Miska and Ferguson-Smith, 2016***; ***Cavalli and Heard, 2019; Sarkies, 2020; Ashe et al., 2021)*.**

The persistence of epigenetically acquired traits in animal populations can be however extremely variable, lasting from a few number of generations to forever. Epigenetic regulations of genes expressed plastically in responses to environmental challenges may become indeed fixed and stably inherited in a given state of expression. This can result in the constitutive expression of a previously plastic state (***Waddington, 1961; Adrian-Kalchhauser et al., 2020***).

Such genetic assimilation may be however a major mechanism through which evolve the innately expressed animal behaviors, traditions or cultures (***Avital and Jablonka, 2000***; ***Crispo, 2007; Renn and Schumer, 2013***). Culture can be defined as a collective adoption and transmission of different behaviors among a sub-population. Until recently, cultural diversity were thought to be restricted to highly cognitive species, as primates and birds. It is now admitted that culture also exist in a great variety of animal species, including invertebrates (***Bridges and Chittka, 2019; Whiten, 2021)*.**

Innate frames underlie the species-specific abilities to respond to environmental challenges and eventually acquire new behaviors. The scope of phenotypic plasticity could extend either randomly, empirically, or both.

The sensory imprinting behavior is an innately encoded environment-directed species-specific behavior. It illustrates the vital necessity of co-incidence between encoded innate frames and actual sensory environment young animals come into the world (***Bateson, 1966; Lorenz and Martin, 1971***).

A high level of phenotypic plasticity, allowing variable developmental and behavioral responses to changing environment, may explain the world-wide expansion of the *C. elegans* nematode species. Worms evaluate their external world mainly through the use of a highly developed chemosensory sense to locate food and avoid toxic chemicals (***Bargmann et al., 1993; Troemel et al, 1995; Bargmann, 2006***). The *C. elegans* chemo-responsive behavior is moreover greatly inluenced by the external environment ***(Remy and Hobert, 2005; Rechavi et al., 2014; Jin et al., 2016; Hong et al., 2017; Posner et al., 2019; Kaletsky et al., 2020; Fernandes de Abreu et al., 2020)*.**

*C. elegans* worms exposed to attractive odors during the critical first larval stage keep life-term imprints of early olfactory experiences. Imprinting increases odor-specific attraction in adult animals. The imprinting behavior is transmitted from odor-exposed to the next unexposed generation, then erased. It can be also stably fixed in worm populations as a new innate behavior, after five consecutive generations were exposed to the same odor.

As previously demonstrated, the innately expressed and the environment-modulated chemo-attractive responses are mediated by different isomers of the Alanine tRNAs (UGC) (***Fernandes de Abreu, 2020***). One form of Ala tRNA (dev-tRNA), processed by ELPC-3, the catalytic sub-unit 3 of the neuronal Elongator complexe, is needed to implement chemo-attractive responses during worm development. The *elpc-3* mutant worms, being unable to synthesize functional dev-tRNAs, do not respond to attractive odors. Feeding *elpc-3* mutants on purified wild-type Ala-tRNA (UGC) however restore the development of wild-type chemo-attractive responses.

Moreover, early exposed worms synthesize odor-specific forms of Ala tRNA (odor-tRNAs). Naive unexposed fed on purified odor-tRNAs acquire the imprinting behavior of odor-exposed.

Innate and acquired worm chemosensations are intrinsically linked, suggesting common regulatory mechanism. Worms must be odor-responsive to be able to perform imprinting. In addition, early exposed or odor-tRNAs fed worms carrying mutations that impair imprinting definitely loose odor-responsiveness (***Fernandes de Abreu, 2020)***. Being stably inherited, such negative imprinting stably supress odor-specific chemo-attractive responses from the innately expressed repertory of worm populations.

In this work, using worm to worm communication protocols, I demonstrate that the regulatory Ala-tRNAs - dev-tRNA and odor-tRNAs - are naturally released in worms environment. All behavioral alterations resulting from feeding biochemically purified Ala-tRNAs are indeed observed using co-culture and food conditioning experiments. Naive acquire imprinting after they were either co-cultivated together with odor-exposed or fed on bacterial food conditioned by odor-exposed, suggesting odor-tRNAs are only released in response to odor-stimuli. By contrast, the *elpc-3* mutants rescuing dev-tRNAs seem constutively secreted by *C. elegans* nematodes.

It is shown here that imprinting and its inheritance require the activity of a number of RNA interference (RNAi) genes. Worm to worm transmission of imprinting is through the systemic RNAi intestinal SID-2 protein (***Winston et al., 2007; McEwan et al., 2012***). It needs at least the first step of the exogenous RNAi pathway, by which externally provided gene-complementary dsRNA triggers long-term transgenerational gene-specific silencing in *C. elegans*. Exogenous dsRNA is processed into small primary siRNAs by a complex associating the Dicer enzyme to the RDE-1 Argonaute, the DRH-1/2 helicases and the dsRNA binding RDE-4 protein (***Timmons et al., 2001; Tabara et al., 2002; Vastenhouw et al., 2006***). Further steps will lead to the synthesis of secondary 26G siRNAs produced by the RdRP RRF-3 and the RNAse ERI-1 *(**Allison et al., 2006;*** ***Han et al., 2009***).

Imprints are stably assimilated as soon as the first generation in *eri-1 (mg366)* mutant worms lacking a functional ERI-1 (***Kennedy, 2004; Thomas, 2014***). Crossing experiments indeed suggest a role for the male ERI-1 in the negative regulation of imprinting inheritance.

Olfactory imprinting in *C. elegans* appears as a deterministic process, part of an obligatory developmental program, based on erasable or heritable epigenetic modifications affecting the expression levels of specific genes in particular cells at a particular stage of development (***Jablonka and Lamb, 2015***).

Positive and negative imprinting respectively increase or decrease the attractive value of an odorant. Imprinting inheritance is stable enough to be analysed genetically. Using genetic crosses, it is shown that attractive values for a single odorant can segregate into five different stable behavioral phenotypes. Both positive and negative imprints behave as gene-specific monoallelic modifications. Crossing imprinted worms generate three behavioral phenotypes following Mendelian segregation rules with 1/4 naive « homozygotes », 1/2 « heterozygotes », and 1/4 « homozygotes » carrying two imprints with additive effects on chemo-attractive responses. Importantly, behavioral « heterozygosity » is stably maintained in hermaphrodite self-fertilized worms.

Crossing worms carrying stable positive imprints for two different odors confirmed typical Mendelian segregation patterns predicted when crossing two gene-specific monoallelic variants. Such crossing indeed generates nine worm populations, each expressing a unique combination of chemo-sensory behavior. A model is proposed in which secreted Ala tRNAs processed by the RNAi machinery, would produce small siRNAs potentially able to interact with the Extra-TFIIIC (ETC) chromosomal locations. They may either maintain or modify chemo-sensory receptor genes expression levels, via unstable or stable chromatin structural changes, according to the chemosensory environment.

## RESULTS

### Worm-to-worm transmission of early olfactory experiences

*C. elegans* L1 larvae imprinting of chemo-attractive odorant molecules enhances odor-specific responses of adults. The imprinting behavior is vertically inherited, either transiently by a single unexposed generation, then erased, or stably after five worm generations were odor-exposed to the same cue (***Remy, 2010***). A previous study showed that odor-stimulated worms produce odor-specific isomers of Alanine tRNA with the UGC anticodon, called odor-tRNAs (***Fernandes De Abreu, 2020***). Naive animals fed on odor-tRNAs purified from worms exposed to three different attractive odors - benzaldehyde (BA), citronellol (CI) and isoamyl alcohol (IA) - acquire the same odor-specific heritable behavioral alterations as if they were odor-exposed. A single Alanine tRNA (UGC) molecule could thus be chemically modified as to carry odor-specific codes, according to worms early olfactory experiences.

Moreover, naive unexposed animals produce a form of Ala tRNA (UGC) which is required to implement the chemo-attractive behavior. Biosynthesis of this « developmental » Ala tRNA requires the activity of ELPC-3, the catalytic sub-unit 3 of the tRNA modifying Elongator complex (***Fernandes De Abreu, 2020***). Inactivating mutations in the *elpc-3* gene impairs the development of chemo-attractive responses, while feeding *elpc-3* mutants on Ala tRNA (UGC) purified from naive wild-type fully rescues chemo-attraction. Conversely, wild-type fed on Ala tRNA (UGC) purified from *elpc-3* mutants acquire the mutant behavioral phenotype.

These results ascribed an essential regulatory role to Ala tRNAs isomers as mediators of the *C. elegans* chemoattractive behavior. However, they were acquired using experimental addition of biochemically purified Ala tRNAs to worm food.

I therefore asked if the same regulations could be observed in more natural conditions, that is without any addition of purified Ala tRNA molecules to worms environment.

To see if odor-exposed worms communicate the imprinting behavior to naive sharing the same environment, I first performed co-culture experiments. As schematically described in **Figure 1A**, wild-type worms carrying an YFP reporter (N2 syls179 (lin-4::yfp) were exposed to the attractive citronellol dilution 1/300 during 24 hours at 20°C from egg laying, a period which encompasses the L1 larval imprinting critical stage (***Remy and Hobert, 2005***). Synchronized wild-type N2 were kept naive unexposed. Part of the YFP-labelled CI-exposed* larvae were mixed together with part of naive larvae (Co-culture), while the other parts of both populations were cultured alone. After 72 hours at 20°C, YFP CI-exposed* were separated from unlabelled naive using a fluorescence microscope. Four adult worms populations were obtained and submitted to CI 300 chemotaxis assays (Figure 1B). As expected, the CI-exposed YFP worm populations 1 and 2 behave as CI-imprinted and migrate significantly faster than the naive unexposed population 4 in a CI 300 gradient. However, worms 3 co-cultured together with CI-exposed worms 2, although naive unexposed, behave as CI-imprinted (**Figure 1B**). This result indicates that naive animals grown together with odor-exposed in the same culture dish adopt the behavior of odor-exposed.

**Figure 1:**
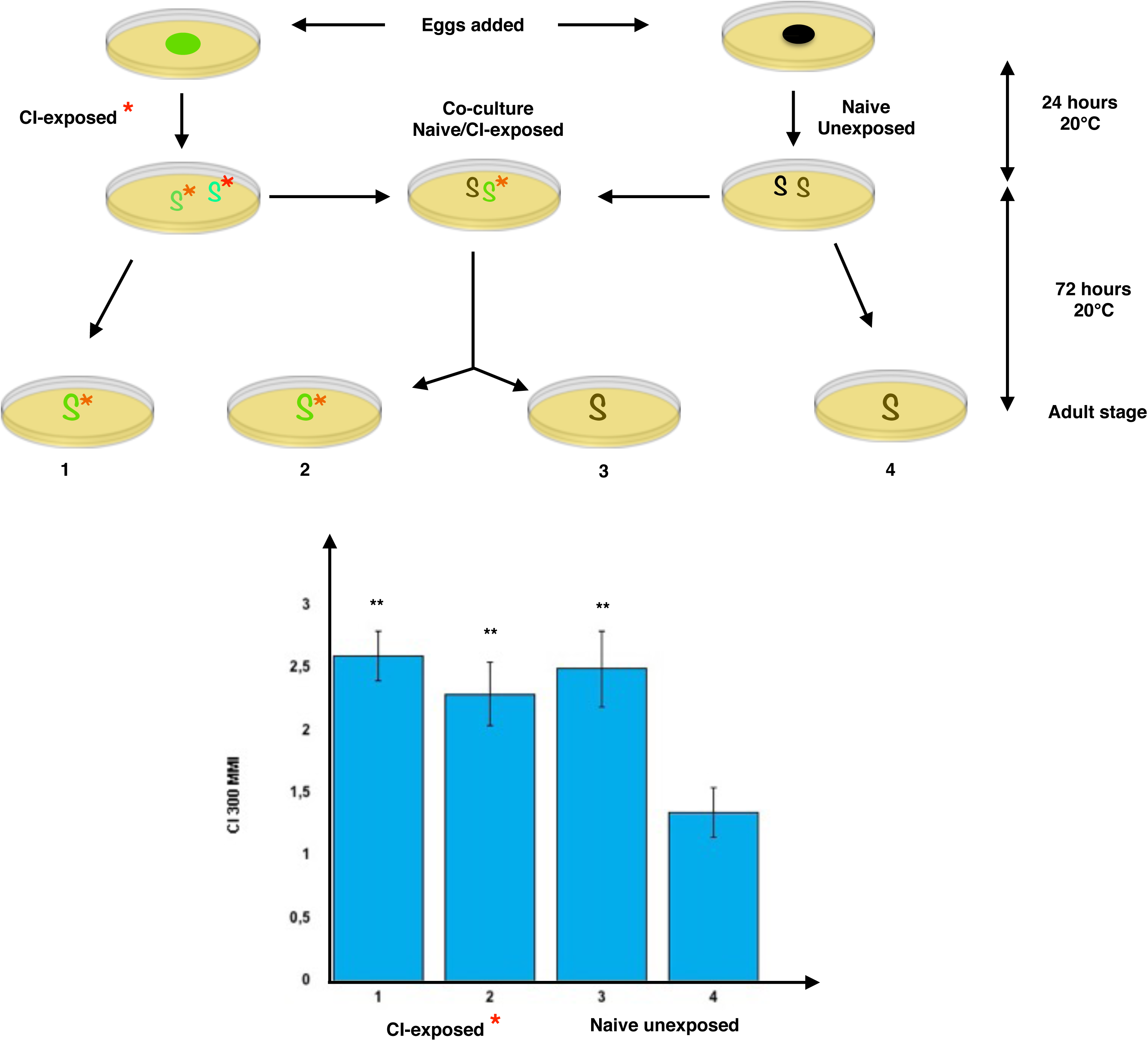
Worm to worm communication of early olfactory experiences. A) Black worms (S) are wild-type N2 *C. elegans*. Green worms (S) are wild-type N2 expressing the YFP fluorescent protein under the control of the *lin-4* promoter (N2 syls179 (lin-4::yfp). YFP worms were exposed (S*) to a chemo-attractive dilution of citronellol (CI 1/300) during 24 hours at 20°C from the egg stage (CI-exposed*). Synchronized wild-type N2 worms were naive unexposed (S). 24 hours old odor-exposed and naive larvae were transferred to new culture plates, either separately (S*S* and SS) or mixed together (SS*). After 72 hours at 20°C, worms reached the adult stage. Cocultured naive and odor-exposed were separated. The four worm populations (Odor-exposed 1 and 2, Naive 3 and 4) were submitted to comparative chemotaxis assays. B) Population chemotaxis assays were performed as described in Methods. Chemotaxis assays determine Mean Migration Indices (MMI) values that account for the mean speed at which an adult worm population migrates up a gradient of an attractive odorant. The CI 1/300 MMI of adult odor-exposed (1 and 2, CI-exposed*) is significantly higher than the CI 1/300 MMI of adult Naive unexposed cultured alone (4). However, Naive unexposed (3) that have been co-cultivated together with CI-exposed (2), display the CI 1/300 MMI of the CI-exposed 1 and 2 worms.

Same results have been obtained using two other chemo-attractive dilutions of benzaldehyde 1/300 and isoamyl-alcohol 1/300 for which early exposure was shown to trigger the production of odor-specific Ala tRNAs (***Fernandes De Abreu, 2020***).

### Horizontal transmission of imprinting requires the intestinal dsRNA transporter SID-2

It was shown that the *sid-2 (qt13)* mutants that do not express the double-stranded selective transporter SID-2 (SID for systemic interference deficient) are unable to acquire olfactory imprinting via feeding purified odor-tRNAs (***Fernandes De Abreu, 2020***). SID-2 is exclusively localized to the apical membrane of intestinal cells, where it is responsible for binding and internalization of large dsRNA from the intestinal lumen. By contrast, absence of the neuronally expressed dsRNA selective importer SID-1 in *sid-1 (qt2)* mutants impairs imprinting inheritance, but not odor-tRNA mediated imprinting transfer (***Fernandes De Abreu, 2020***).

Co-culture experiments were performed as described in **Figure 1A**. CI 200 was used instead of CI 300 imprinting since *sid-2 (qt13)* worms migrate slowly than wild-type toward all chemo-attractants. As shown in **Figure 2**, naive *sid-2 (qt13)* MMI to CI 200 remains unaffected, even after life-long co-culture together with naive or with CI 200 exposed N2. As a control, the same experiment was performed using *sid-1 (qt2)* / N2 co-cultures. Co-culture with naive N2 does not modify the *sid-1 (qt2)* MMI, while co-culture with CI 200 exposed N2 significantly increases the *sid-1 (qt2)* MMI to CI 200.

**Figure 2:**
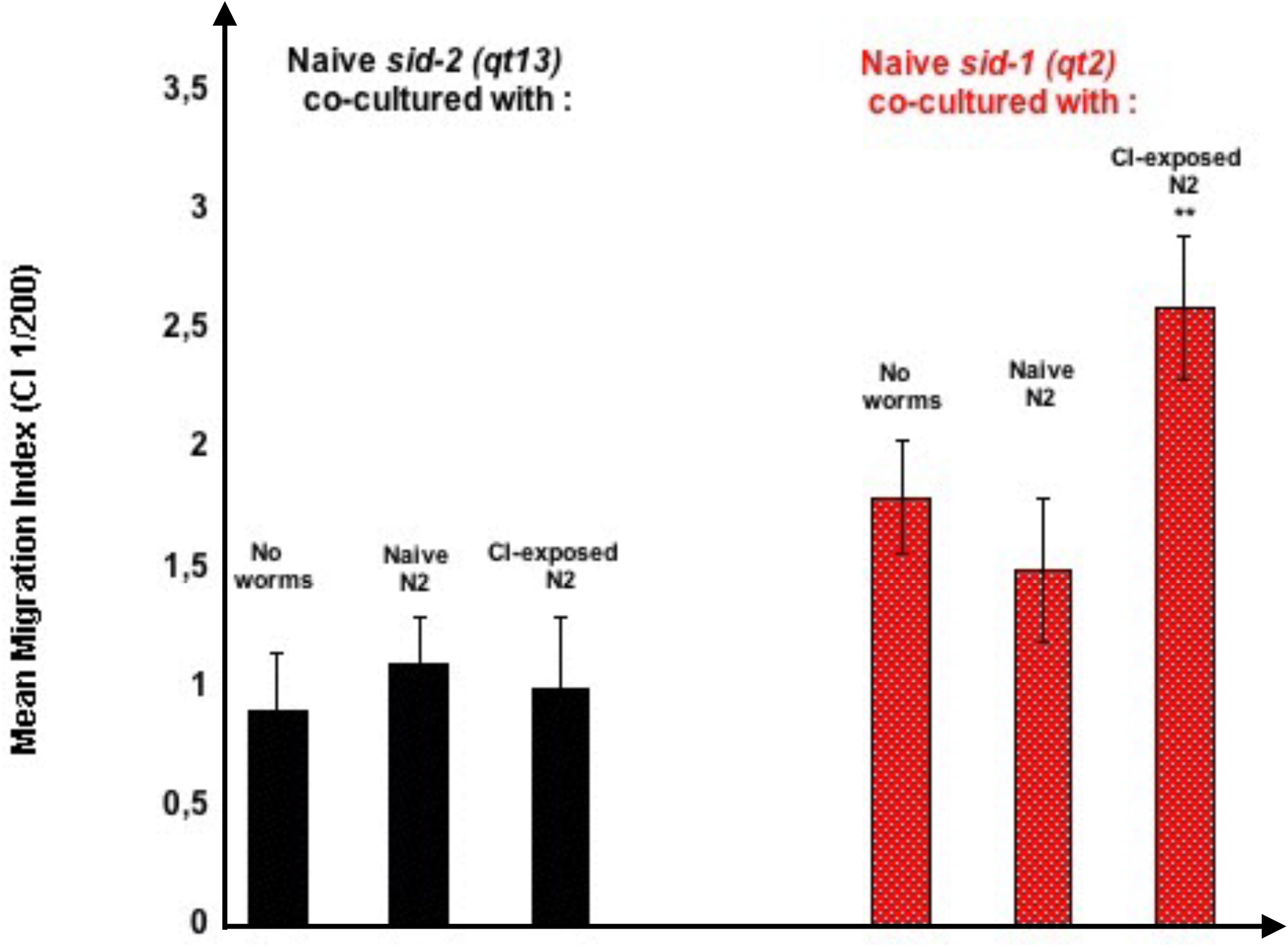
The horizontal transmission of imprinting requires the systemic RNA interference (RNAi) protein SID-2. Co-culture experiments were performed as described in Figure 1. Black bars: Co-culture with CI-exposed N2 does not modify the Mean Migration of *sid-2 (qt13)* worms in a CI 1/200 gradient, compared to culture alone (No worms) or co-culture with naive wild-type (Naive N2). Red bars: Co-culture with CI-exposed N2 significantly increase the Mean Migration of *sid-1 (qt2)* worms in a CI 1/200 gradient, compared to culture alone (No worms) or co-culture with naive wild-type (Naive N2).

Inter-individual communication of a behavioral phenotype involve the production and emission of significant signals by animals that express the phenotype. Being received by others, these signals influence their behavior. From **Figure 2** data, it can be concluded that, since none RNA molecules but odor-tRNAs specifically transfer olfactory imprinting from exposed to naive, CI-exposed worms release CI-specific Alanine tRNA (UGC) in the external environment.

Secreted odor-tRNAs might be processed after entering worm bodies through SID-2, shown to be the entry point for externally provided dsRNAs (***Winston et al., 2007; McEwan et al., 2012; Marré et al., 2016***). How naive further read and translate the odor-tRNAs messages into behavioral changes is unknown. The horizontal transmission of imprinting by external tRNAs entering naive worm tissues through SID-2 however suggest that an early step of the imprinting process may involve processing by the exogenous RNA interference (RNAi) pathway.

### Exogenous RNAi controls olfactory imprinting

Gene silencing by RNA interference (RNAi) was discovered in *C. elegans*. RNAi has been and is still widely used experimentally to knock down gene expression via double-stranded RNAs homologous to endogenous sequences. In *C. elegans*, gene silencing can be achieved via injection or via feeding worms with artificially-expressed dsRNAs (***Timmons and Fire, 1998; Tabara et al., 1998; Timmons et al., 2001***).

Using genetic screens, a number of mutant worms resistant to gene silencing after exogenous addition of dsRNA had been isolated. The first step of the exogenous RNAi pathway is dsRNA processing into short interfering primary siRNAs by the Dicer complex. This complexe associates the RNase III Dicer DCR-1, an Argonaute RDE-1, two Dicer-related helicases DRH-1 and 2 and the double-stranded RNA binding protein RDE-4 (***Tabara et al., 2002***).

Some other RNAi-resistant mutations, as *rde-2/mut-8* and *mut-7 (pk204),* were found to inactivate the mutator complex, also involved in transposon silencing (***Ketting et al., 1999***).

Gene silencing by RNAi is trans-generationally transmitted (***Vastenhouw et al., 2006***) and systemic in *C. elegans*, as it spreads across worm tissues using the different SID (<u>s</u>ystemic interference <u>d</u>eficient) proteins (***Timmons et al., 2003***).

*C. elegans* worms express RNA-dependent RNA polymerases (RdRPs), as RRF-1 and RRF-3, required for the amplification and trans-generational transmission of gene silencing through the synthesis of 22G and 26G secondary siRNAs. Worms carrying mutations inactivating RRF-1 as *rrf-1 (pk1417)* and *rrf-1 (ok589)* are RNAi resistant.

A number of mutations have been isolated that increase the efficiency of exogenous RNAi (<u>e</u>nhanced <u>R</u>NA interference mutants). Among them, *rrf-3 (1426)* inactivates the RRF-3 RdRP, while *eri-1 (mg366)* impairs the function of the 3’ exonuclease ERI-1. Both RRF-3 and ERI-1 belong to the endogenous RNAi pathway, and control the endogenous production of 26G long secondary siRNAs (***Allison et al., 2006; Sijen et al., 2007; Gu et al., 2009; Han et al., 2009; Gent et al., 2010; Xu et al., 2018***).

The effects of RNAi resistant and RNAi enhancing mutations on imprinting and its inheritance were assessed (***Figure 3***). In order to represent the behavioral phenotype of RNAi mutants by a single value, results are expressed in Mean Imprinting Indices (MII). The MII values are obtained after substraction of the Mean Migration Index (MMI) value of naive from the MMI value of exposed. A positive MII value thus indicates exposed worms migrate faster that naive in chemotaxis assays, reflecting a positive imprinting, as in N2 wild-type. Conversely, a negative value for MII indicates exposed worms migrate slower than naive toward the odor source, reflecting a negative imprinting.

**Figure 3:**
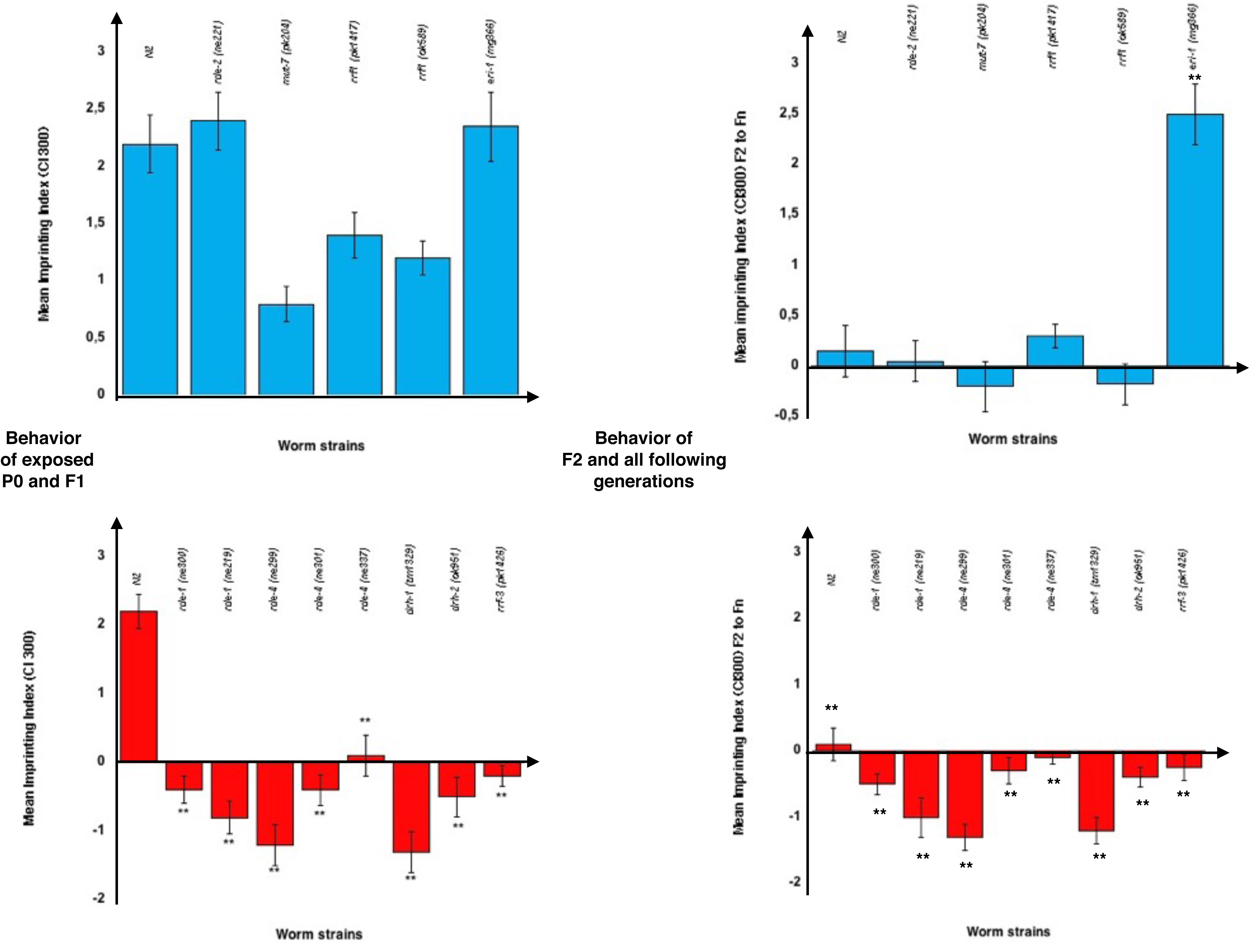
RNA interference (RNAi) genes control imprinting and its inheritance. **Figure 3A : Positive and Negative olfactory Imprinting in RNAi mutants** Mean Imprinting Indices (MII) represent the mean difference between the Mean Migration Index (MMI) of odor-exposed worms and the MMI of unexposed naive worms. Positive MMI values are observed when the CI 1/300 exposed worms migrate significantly faster than the naive unexposed worms. Negative MMI values are observed when CI 1/300 exposed worms migrate more slowly than naive unexposed worms. Blue bars: Positively imprinted RNAi mutants Positive Mean Imprinting Indices are observed for the *rde-2 (ne221), mut-7 (pk204), rrf-1 (pk1417), rrf-1 (ok589)* and *eri-1 (mg366)* mutant strains. The N2 MMI value and the RNAi mutants MMI values for CI 1/300 are, respectively, 2.2 ± 0.3, 2.4 ± 0.25, 0.7 ± 0.15, 1.23 ± 0.06, 1.1 ± 0.08 and 2.35 ± 0.3. Red bars: Negatively imprinted RNAi mutants Negative Mean Imprinting Indices are observed for the *rde-1 (ne300), rde-1 (ne219), rde-4 (ne299), rde-4 (ne301), rde-4 (ne337), drh-1 (tm1329)*, *drh-2 (ok951)* and for *rrf-3 (pk1426)* mutant strains. All mutants MMI values for CI 1/300 are significantly lower (** p<0.01) than the N2 MII value of 2.2 ± 0.3, with, respectively, -0.4 ± 0.2, -1 ± 0.24, -1.45 ± 0.3, -0.45 ± 0.23, 0.15 ± 0.3, -1.2 ± 0.27, -0.42 ± 0.29, and -0.2 ± 0.3. **Figure 3B : Stable or unstable inheritance of olfactory imprints** Blue bars: Positive imprinting is erased beyond the F1 generation in N2 and in all *rde-2 (ne221), mut-7 (pk204), rrf-1 (pk1417), rrf-1 (ok589)* RNAi mutants. It is by contrast stably transferred in the progeny of exposed worms carrying the *eri-1 (mg366)* mutation. Red bars: Negative MII imprinting values are stably transmitted in the progeny of the *rde-1 (ne300), rde-1 (ne219), rde-4 (ne299), rde-4 (ne301), rde-4 (ne337), drh-1 (tm1329)*, *drh-2 (ok951)* and *rrf-3 (pk1426)* RNAi mutants.

Some mutations inactivating or enhancing exogenous RNAi have no effect on the imprinting behavior. A wild-type behavior, assessed by a positive MII values for CI 300 imprinting, is observed for worms carrying the RNAi resistant *rde-2 (ne221), mut-7 (pk204), rrf-1 (pk1417), rrf-1 (ok589)* mutations, and for the enhanced RNAi mutant *eri-1 (mg366)* (***Figure 3A, Blue bars***).

All independent alleles inactivating members of the primary siRNAs producing Dicer/RDE-1/DRH-1/2/RDE-4 complex, respectively *rde-1 (ne300), rde-1 (ne219)*, *drh-1 (tm1329), drh-2 (ok951), rde-4 (ne299)*, *rde-4 (ne337)* and *rde-4 (ne337)*, were found however unable to produce a positive imprint of CI 300 (***Figure 3A , Red bars***). As a result, the Mean Imprinting Index values are, thus to different extents, negative for these mutants. Instead of promoting an enhanced response to the imprinted odors, as in N2, the output of early odor-exposure in these mutants is to decrease adult responses, compared to naive. It has to be noted that worms carrying *sra-11*, as well as *elpc-1* mutations, previously described as olfactory imprinting deficient, were also stably desensitized after odor-exposure (***Remy and Hobert, 2005; Fernandes De Abreu et al., 2020.***

With the notable exception of the *eri-1 (mg366)* mutants, positive imprinting is transmitted to the F1 generation, then erased (***Figure 3B, Blue bars***). By contrast, negatively imprinted worms, as already shown for the *elpc-1* mutants, stably transfer their behavior to all generations following the odor-exposed (***Figure 3B, Red bars***).

### Alanine tRNA (UGC) are released in worms environment

Laboratory *C. elegans* worms are routinely cultured on agar plates loaded with slow-growing *Escherichia coli* (OP50 strain) lawns as a food source.

I used the food-conditioning protocol schematically described in **Figure 4** to demonstrate that both naive and odor-exposed worms release different forms of Alanine tRNAs (UGC) in their environment.

**Figure 4:**
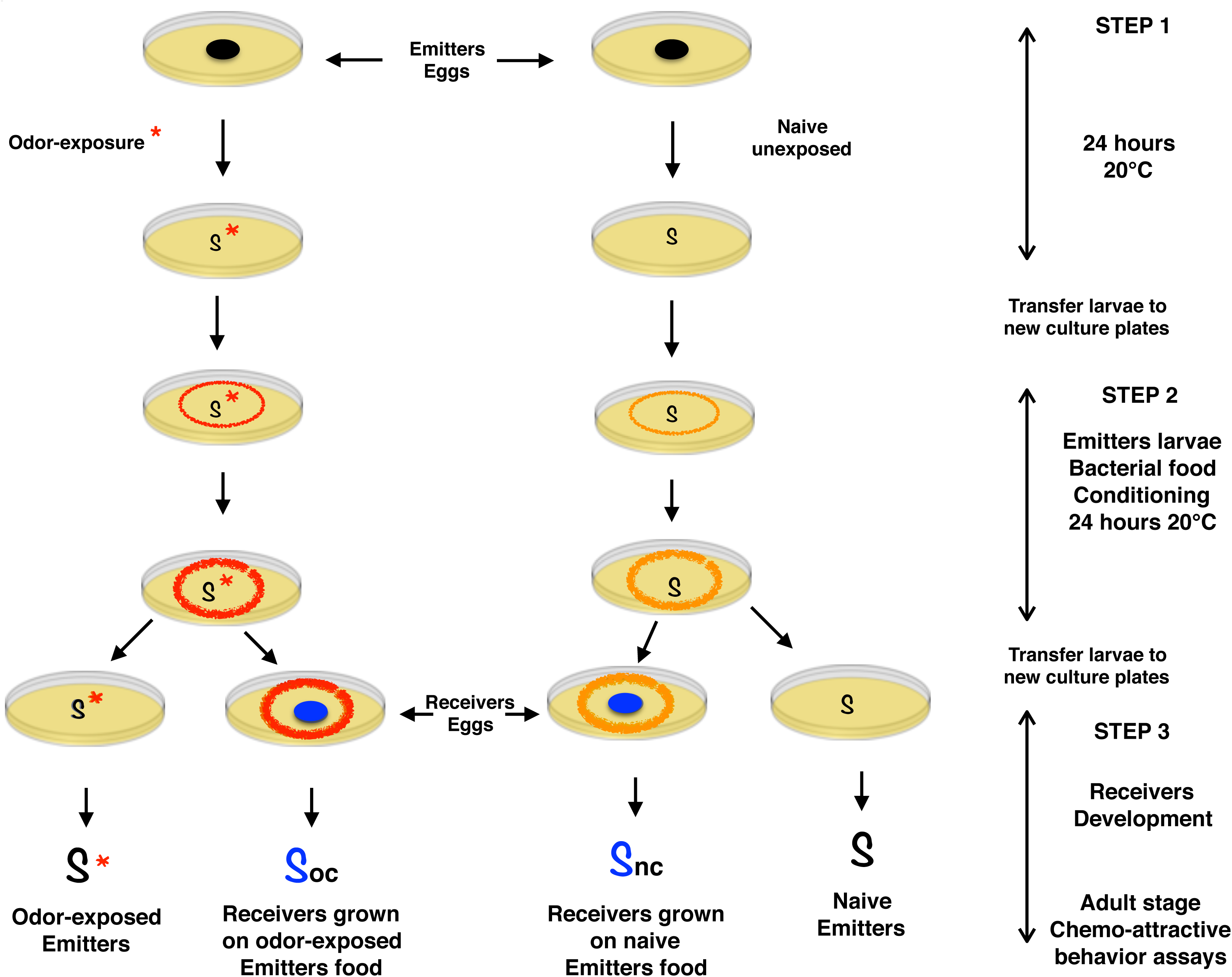
Conditioned bacterial food by odor-experienced or naive worms. Culture plates are loaded with bacterial spots (OP50 strain of E. coli, the standard worm food) of approximately one square centimeter. Emitters (black) and Receivers (blue) worms belong either to the same or to different genetic strains. STEP 1 : Emitters are - or are not for naive controls - odor-exposed during the critical imprinting period of development uncovering the first larval stage L1 (24 hours at 20°C from the egg stage). Odor-exposed or naive Emitter larvae are transferred to new culture plates. STEP 2 : The progressive conditioning of bacterial food due to the presence of worm larvae during variable time periods is illustrated by, respectively, orange rings for naive or red rings for odor-exposed (Conditioning). At the end of STEP 2, Emitters larvae are transferred to new plates to resume development up to the adult stage. STEP 3 : The whole egg to adult development of 20 Receivers worms takes place on bacterial food conditioned either by naive or by odor-exposed Emitters. The chemo-attractive responses of Odor-exposed or Naive Emitters, of Receivers grown on Odor-exposed or on Naive Emitters conditioned food, were compared at the adult stage, as described in Material and Methods.

Emitters are either odor-exposed (**S**\*) during 24h at 20°C or kept naive (**S**) unexposed (STEP 1). Emitter larvae are then transferred to fresh culture plates loaded with equal amounts of bacterial food (about 3 cm^2^ spots) and grown during 24 hours at 20°C (STEP 2). During the food-conditioning STEP 2, odor-exposed or naive growing larvae may or may not leave their imprints onto the bacterial spots. Food conditioning by odor-exposed worms are represented by red circles, while orange circles represent food conditioning by naive animals.

Using exposed N2 as Emitters and naive N2 as Receivers, I determined that imprinting transfer via feeding requires food has been conditioned by the presence of at least five 24 hours old Emitter larvae during 24 hours at 20°C.

48 hours old Emitter larvae - a developmental stage at which worms do not yet lay eggs - were again transferred at the end of STEP 2. Receivers (**S**) are always naive unexposed. Receivers are growing from egg to adult stages on bacterial lawns conditioned by the presence of either odor-exposed (**Soc** for <u>o</u>dor-exposed <u>c</u>onditioned) or of naive (**Snc** for <u>n</u>aive <u>c</u>onditioned) emitter larvae (STEP 3). Comparative chemotaxis assays were performed at the adult stage.

Emitter and Receiver worms can belong either to the same or to genetically different strains. Comparing the behavior of odor-exposed (**S**\*) to naive (**S**) Emitters indicates if and how early odor-exposure has been imprinted, while comparing the behavior of their progeny informs on imprinting inheritance.

Emitter mutants - i.e. worms that do not release the imprints - are identified by comparing the behavior of (**Soc**) to the behavior of (**Snc**), when wild-type N2 is used as the Receiver strain.

Comparing the behavior of **Soc** to the behavior of **Snc** identify Receiver mutants - i.e. worms that are not imprinted by the release of imprints -, when wild-type N2 is used as the Emitters strain.

Results are summarized in **Table 1:**

**Table 1:** Odor-tRNAs and dev-tRNAs are released in worms environment. Column 1: Odor-exposed (CI 300 or BA 300) worms carrying the indicated mutations were the Emitter worms, as described in Figure 4. Naive N2 were the Receivers. To determine if imprinting was (**+**) or was not (**-**) transferred, Receiver worms were submitted to chemotaxis assays. Column 2: Receiver worms carrying the indicated mutations were fed on bacterial food conditioned by odor-exposed N2. The behavioral effects triggered by feeding N2 released imprints are described. Column 3: Naive mutants were used as Emitters. Receivers were naive chemo-attraction deficient *elpc-3 (ok2452)* or naive *elpc-3 (tm3120).* Chemotaxis assays determined if the chemo-attractive responses of the *elpc-3* mutants were (**+**) or were not (**-**) restored.

|  | Exposed mutants<br>as Emitters<br>*Receivers N2* | Exposed N2 as Emitters<br>*Receivers mutants*<br>*Behavioral changes* | Naive mutants as<br>Emitters<br>*Receivers elpc-3* |
| --- | --- | --- | --- |
| Chemotaxis |  |  |  |
| *odr-1 (n1936)*<br>*tax-2 (p671)*<br>*tax-4 (p678)* | - | Unchanged | + |
| Exogenous RNAi |  |  |  |
| *rde-2 (ne221),*<br>*mut-7 (pk204)*<br>*rrf-1 (pk1417)*<br>*rrf-1 (ok589)* | + | Unstable positive imprinting | + |
| *eri-1 (mg366)* | + | Stable positive imprinting | + |
| *rde-1 (ne300)*<br>*rde-1 (ne219)*<br>*rde-4 (ne299)*<br>*rde-4 (ne301)*<br>*rde-4 (ne337),*<br>*drh-1 (tm1329),*<br>*drh-2 (ok951)*<br>*rrf-3 (pk1426)* | + | Stable negative imprinting | + |
| Negatively<br>Imprinted F1 to Fn<br>*rde-1, rde-4, drh-1,*<br>*drh-2, rrf-3*<br>*mutants* | - | Unchanged | + |
| Systemic RNAi |  |  |  |
| *sid-1 (qt2)* | + | Unstable positive imprinting | + |
| *sid-2 (qt13)* | + | Unchanged | + |
| Elongator<br>complex |  |  |  |
| *elpc-1 (ng10)* | + | Stable negative imprinting | + |
| Negatively<br>Imprinted F1 to Fn<br>*elpc-1 (ng10)* | - | Unchanged | + |
| *elpc-3 (tm3120)*<br>*elpc-3 (ok2452)* | - | Unchanged | - |
| Ascaroside<br>biosynthesis |  |  |  |
| *daf-22 (m130)* | + | Unstable positive imprinting | + |

**First column:** The indicated strains were used as odor-exposed Emitters, while Receivers were naive. According to whether exposed Emitters do (+) or do not (-) release odor-tRNAs imprints, N2 Receivers will or will not be positively imprinted (as shown in **Figure 1**).

**Second column**: Exposed N2 were used as odor-tRNAs Emitters. Receivers carry the indicated mutations. The Receivers behavioral changes resulting from feeding food conditioned by exposed N2 is described.

**Third Column**: The indicated worm strains were kept unexposed naive and used as Emitter. Receivers were the chemoattraction deficient *elpc-3* mutants. If Emitters release the « developmental » form of wild-type Ala tRNA, chemo-atttractive responses will be fully rescued in *elpc-3* mutants used as Receivers.

The *odr-1 (n1936)*, *tax-2 (p671)* and *tax-4 (p678)* mutations impair the transduction of olfactory messages downstream the interaction of odorant molecules with putative olfactory G-protein coupled receptors (GPCR) expressed by the chemo-sensory AWC neurons (***Bargmann et al., 1993; Bargmann, 1996***). These chemotaxis mutants do not synthesize nor release odor-tRNAs imprints (-) upon odor-exposure. They, however, produce and secrete the *elpc-3* rescuing developmental form of Ala tRNA.

Exogenous RNAi resistant mutants are odor-responsive and do release odor-tRNAs after odor exposure (+). Worms with mutations inactivating the *rde-2*, *mut-7*, *rrf-1* and *eri-1* genes acquire a positive imprinting after feeding odor-tRNAs released by exposed N2. Worms with mutations in the *rde-1*, *rde-4, drh-1/2* and *rrf-3* genes stably imprint via feeding odor-tRNAs, however negatively (**Figure 3**). Negatively imprinted worms have lost odor receptivity: as a result, they do not longer release odor-tRNAs in the medium when odor-exposed (-).

However, all tested RNAi mutant worms, even those that were negatively imprinted, do release the *elpc-3* rescuing developmental Ala tRNA (+).

The *sid-1 (qt2)* mutants behave as wild-type: they are odor-tRNA Emitters after odor-exposure (+), developmental Ala-tRNA Emitters (+), and positively imprinted when used as odor-tRNAs Receivers. By contrast, the *sid-2 (qt13)* mutants, as already shown on **Figure 2**, do not receive the imprinting odor-tRNA messages send by exposed N2 Emitters. Importantly, *sid-2 (qt13)* worms do respond to chemo-attractive odorants, however with lower mean migration indices than N2. Nevertheless, the *sid-2 (qt13)* mutants positively imprint chemo-attractants after early odor exposure.

Both systemic RNAi *sid-1* and *sid-2* mutants do release the developmental Ala-tRNA while naive (+).

Exposed *elpc-1 (ng10)* mutants are odor-tRNAs Emitters (+), while they imprint negatively after feeding exposed N2 conditioned food. Negatively imprinted *elpc-1 (ng10)* worms lost odor responsiveness, thus no longer produce nor release odor-tRNAs upon odor-exposure (-). Naive unexposed as well as negatively imprinted *elpc-1 (ng10)* worms however secrete the developmental form of Ala tRNA (+).

*elpc-3 (tm3120)* and *elpc-3 (ok2452)* do not develope a chemo-attractive behavior. The Elongator complex catalytic sub-unit 3, ELPC-3, regulates the chemical modifications pattern of tRNAs bases. The two *elpc-3* mutants would then produce and release a defective, non-functional, form of Ala tRNA (UGC), unable to support the development of a wild-type chemo-attractive behavior. Food conditioned by naive worms from all tested genotypes do rescue the chemo-attractive responses in *elpc-3* mutants (Table 1, third column +).

Importantly, worms carrying the ascaroside biosynthesis defective *daf-22 (m130)* mutation behave as wild-type, suggesting ascaroside synthesis and secretion has no influence on the behavioral regulations described in this study.

More detailed data on *elpc-3* rescue are provided by the **Supplemental Figure**: naive N2 were used as Emitters while naive *elpc-3 (tm3120)* and *elpc-3 (ok2452)* worms were the Receivers. Feeding food conditioned by naive N2 restores chemo-attraction up to wild-type levels (MMI to CI 1/300) in both *elpc-3* mutants (Blue bars). The reverse experiment, in which naive *elpc-3* mutants were used as Emitters and naive N2 as Receivers (Red bars), was performed. Feeding N2 on bacteria conditioned by *elpc-3* worms reduced MMI to CI 1/300 down to the mutant levels.

Figure 1 to 4, Supplemental Figure and Table 1 data suggest *C. elegans* nematodes release different forms of regulatory Alanine tRNA in their living environment. The bacterial food conditioned by naive worms from all tested mutant strains, including the chemotaxis and imprinting defective mutants, restore chemo-attraction in *elpc-3* mutants, suggesting the external release of the developmental form of Ala tRNA. Odor stimuli makes all odor-responsive worms release odor-specific Ala tRNAs. They can act as extrinsic signals able to communicate sensory experiences to naive conspecifics. Consistent with a process based on the uptake of environmental RNAs by the intestinal SID-2 protein, imprinting involves at least the first step of the exogenous RNAi pathway, in which dsRNA is processed into primary siRNAs by the Dicer-associated complex members RDE-1, DRH-1/2 and RDE-4 (***Tabara et al., 2002***).

In order to assess the stability of odor-tRNAs while on bacterial lawns, I used Ala tRNA (UGC) highly purified from CI 300 exposed worms (**Supplemental Table**). As already published, the mean purification yield was about 100 pg CI-tRNA per 10^3^ worms (***Fernandes De Abreu et al., 2020***). The indicated increasing amounts of CI-tRNA from 1 fg to 1 ng were deposited on 3 cm^2^ bacterial spots onto which 20 naive N2 eggs were added either immediately (No delay) or after the indicated delays, from 24 to 96 hours at 20°C.

Transfer of a CI imprint to naive N2 receivers was assessed by comparing the CI 300 MMI of CI- tRNA fed to the CI 300 MMI of naive, as described.

As low as 1 fg (10^-15^ g) CI-tRNA is able to imprint 20 naive receivers if eggs and CI-tRNAs were deposited at the same time. As expected, the imprinting transfer efficiency progressively decreased according to the time CI-tRNAs spend on bacterial spots. Deposition of 1 fg tRNA is the minimal amount required to imprint 20 worms. CI imprinting transfer can be observed 24 hours, but not 48 hours, after addition of 10 fg CI-tRNA to the bacterial spots. Deposition of at least 100 pg CI-tRNA is necessary to keep imprinting transfer active after a 48 hours delay, while after 1 ng was deposited, enough active CI-tRNAs - able to transfer CI imprinting - resisted to degradation even after a period of 72 hours on bacterial lawns.

The amount of odor-tRNA molecules entering through intestinal cells, and actually required to trigger a behavioral change in a receiver worm is unknown. However, odor-tRNA mediated horizontal transmission of imprinting seems highly efficient, even with a low number of odor-tRNA molecules present in worms environment.

### The ERI-1 RNAse negatively control genetic assimilation

Stable positive imprinting is achieved in wild-type worms after five generations were odor-exposed or odor-tRNA fed. Remarkably, a stable transmission of positive imprinting is also achieved after a single exposure in the *eri-1 (mg366)* mutants (**Figure 3**).

ERI-1 is a multifunctional protein, bearing a 3’ to 5’ exonuclease RNase activity and a DNA binding SAP domain. The *C. elegans eri-1* gene encodes at least two isoforms of the ERI-1 protein, a 448 amino-acids ERI-1a, and a larger 582 amino-acids ERI-1b. The reference allele *eri-1 (mg366)* is a frame shift variant due to a 23 bp insertion that may affect the synthesis of both ERI-1a and b isoforms.

At the adult stage, ERI-1 is mainly expressed in the spermatheca and in a few neurons ***(Kennedy et al., 2004; Gabel and Ruvkun, 2008; Thomas et al., 2014).*** Due to this expression pattern, it is tempting to hypothezise a « gender-specific » role for ERI-1 in imprinting stability over generations. The main mode of reproduction of hermaphrodite *C. elegans* nematodes is by self-fertilization. Male individuals can be however obtained and used for sexual reproduction, allowing genetic segregation analysis.

To demonstrate ERI-1 prevents the stable assimilation of imprints, three crosses were performed. As shown in **Figure 5**, CI 300 exposed N2 or *eri-1 (mg366)* hermaphrodites were or were not crossed with naive N2 or *eri-1 (mg366)* male individuals. Males carrying YFP or GFP transgenes were used in order to easely distinguish between self (unlabelled) and crossed (YFP or GFP labelled) F1 larvae. Worm populations were generated from individually isolated F1 worms from crossed or self progeny. CI 300 responses were assessed from the F2 generation and beyond. **Figure 5** shows the behavior of the F5 worms. The A cross (exposed N2 X naive N2) was performed as control. As already indicated, imprinting is not transferred beyond F1 in self-fertilized exposed N2 (A Self).

**Figure 5:**
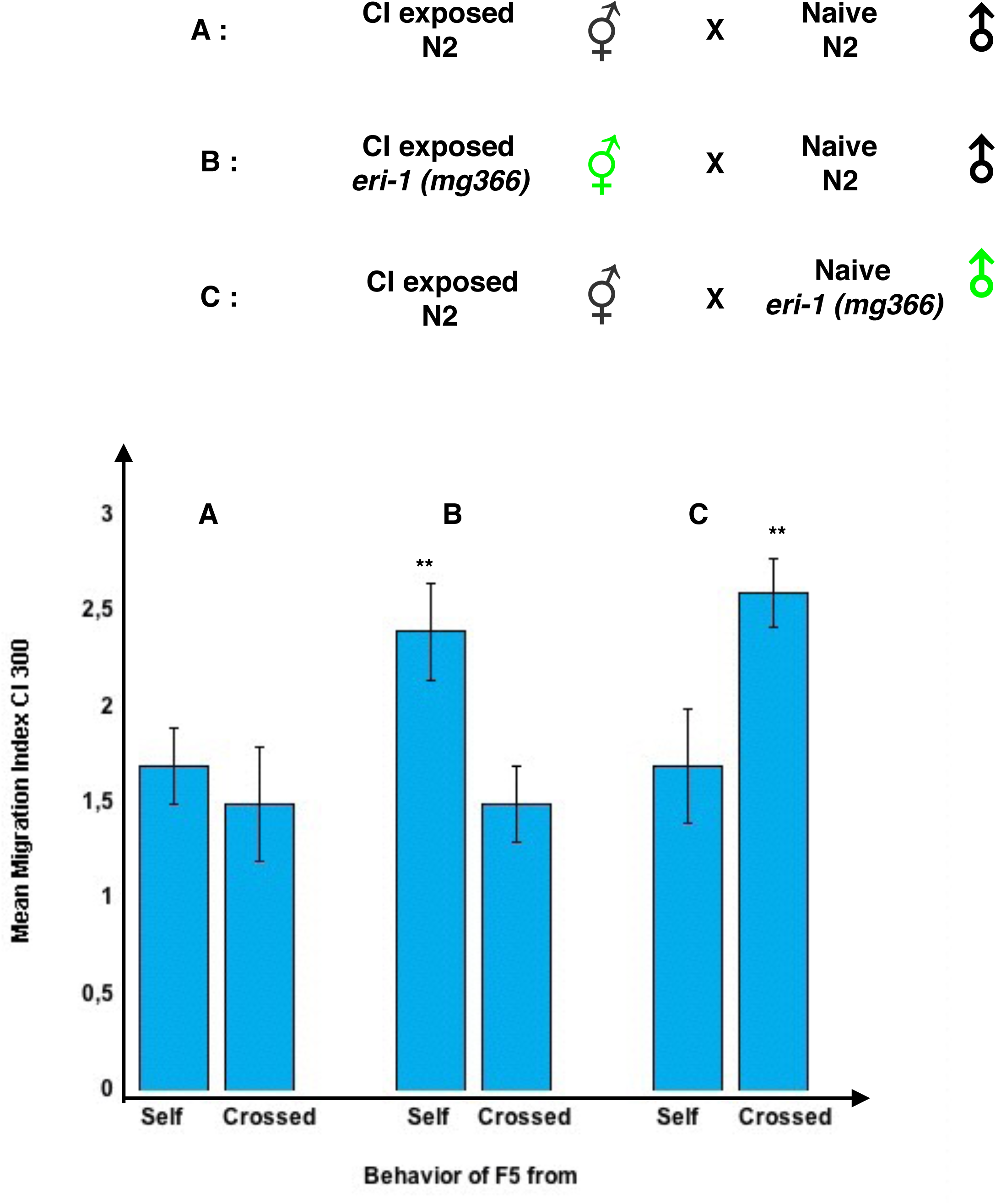
The « paternal » 3’ exonuclease ERI-1 negatively controls the stable inheritance of imprinting. Three crossing experiments were performed: **A**: CI 300 exposed N2 hermaphrodites (⚥) were crossed to naive N2 males (♂), **B**: CI 300 exposed *eri-1 (mg366)* hermaphrodites (⚥) were crossed to naive N2 males (♂), and **C**: CI 300 exposed N2 hermaphrodites (⚥) were crossed to naive *eri-1 (mg366)* (♂) males. Worm populations (n > 10) were generated from F1 individuals, either from self-fertilization (her-maphrodites, Self) or from sexual (hermaphrodites X males, Crossed) reproduction. MMI to CI 300 (Blue bars) are shown for the F5 generation of self and crossed progeny. **A cross**: MMI were as naive for either the Self F5 progeny (A, Self) of exposed N2 or for the Crossed F5 progeny (A, Crossed). **B cross**: MMI were significantly higher (**, p < 0.01, n > 10) for the self F5 progeny of exposed *eri-1 (mg366)* (B, Self) than for the F5 crossed progeny (B, Crossed). **C cross**: MMI were found significantly higher (**, p < 0.01, n > 10) for the crossed F5 progeny (C, Crossed) than for the self F5 progeny (C, Self) of exposed N2.

CI imprints do not persist either in worms issued from exposed N2 crossed with N2 naive males (A Crossed).

Imprinting is stably transmitted in the self-descendance of exposed *eri-1 (mg366)* hermaphrodites (B self). They are however erased in the progeny of exposed *eri-1 (mg366)* hermaphrodites crossed with *eri-1+/+* N2 males (B crossed). By contrast, the imprinting behavior is maintained and stably inherited after exposed N2 were crossed with males that do not express ERI-1 (C Crossed), while, as expected, it does not persist in the self progeny of exposed N2 (C Self). **Figure 5** data suggest the « paternal » 3’-exonuclease ERI-1 activity indeed acts as an eraser. Stable inheritance is achieved after five exposed generations in wild-type. The RNA species responsible for imprinting fixation would accumulate throughout generations, exceeding the ERI-1 erasing potential after five generations.

### Stably inherited positive or negative imprinting behavior segregate as Mendelian monoallelic alterations in the crossed progeny

Positive and negative imprinting produce self-fertilized hermaphrodite worm populations with, respectively, enhanced or decreased chemo-attractive value assigned to the initial odorant trigger. These behavioral changes are odor-induced in all individuals of the exposed generation and are stably maintained in the whole worm progeny. They therefore cannot be due to the selection of pre-existing variations of DNA sequences in worm populations, but might nevertheless be stable enough to be tractable using genetic segregation analysis.

**Figure 6** describes the segregation pattern of chemo-attractive values in the crossed progeny of positively and negatively imprinted worms by either the CI 300, BA 300 or IA 300 attractive odorants. Positively imprinted worms were *eri-1 (mg366)* mutants after a single exposure (odor-stimuli or odor-tRNA feeding), or N2 worms in which imprinting has been stabilized after five consecutive generations were exposed. Negatively imprinted worms were *elpc-1 (ng10)* mutants after a single exposure.

**Figure 6:**
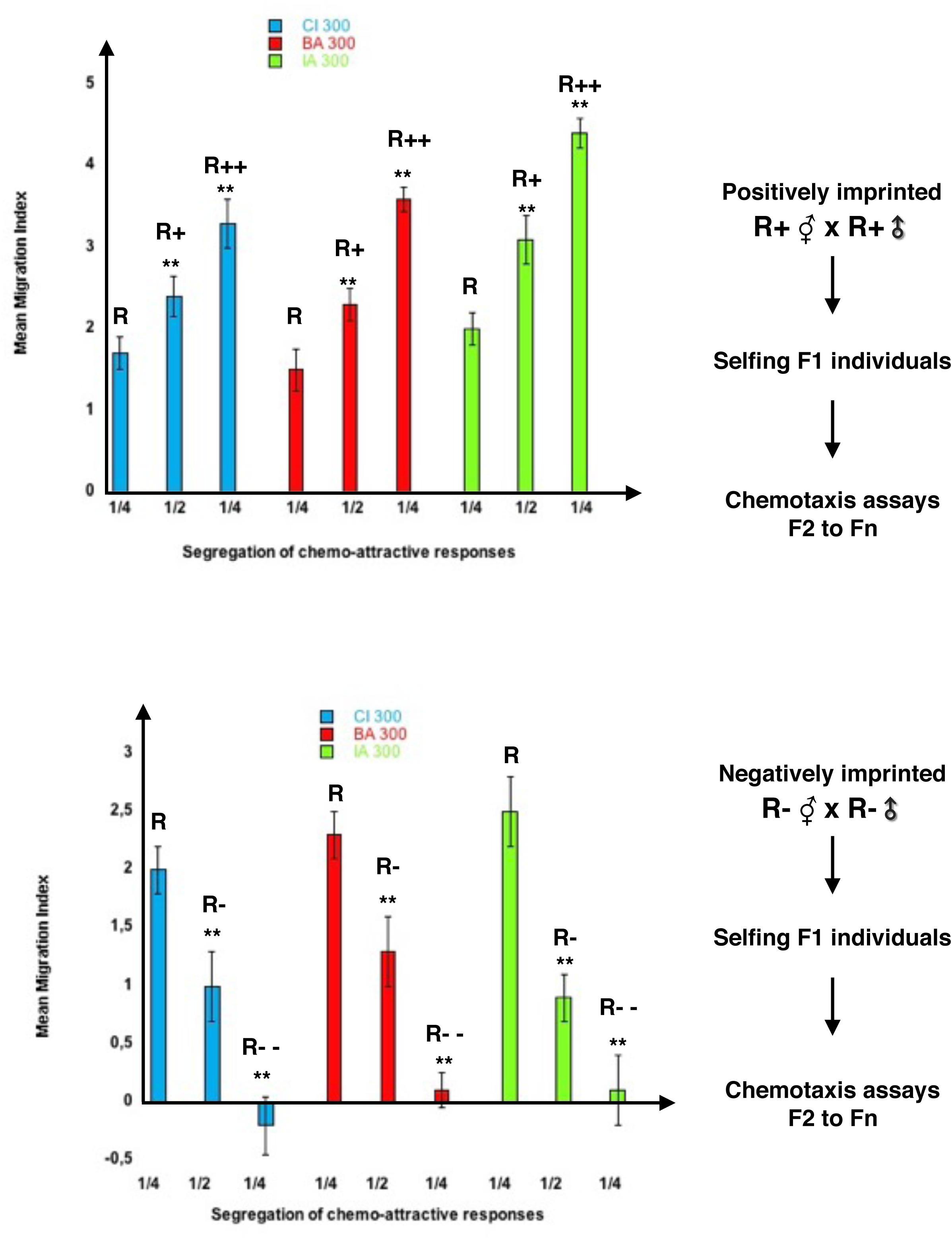
Mendelian segregation of chemo-attractive values in the crossed progeny of stably imprinted worms. 6a) Stably CI 300, BA 300 or IA 300 positively imprinted hermaphrodites (maternal R+) were crossed to their own males (paternal R+). The F2 to F5 progeny of 20 F1 individuals isolated from each R+ x R+ cross were submitted to odor-specific chemotaxis assays. The Mean Migration Indices to the respective imprinted odor dilutions were determined. Chemo-attractive responses segregate into three different stably inherited levels: half crossed worm populations display the maternal behavior (R+), 1/4 lost the positive imprinting and behave as naive (R), while 1/4 show MMI values significantly higher than maternal (R++ > R+ > R, **p < 0.01). 6b) Stably CI 300, BA 300 or IA 300 negatively imprinted hermaphrodites (maternal R-) were crossed to their own males (paternal R-). The F3 to F5 progeny of 20 F1 individuals isolated from each R- x R- cross were submitted to odor-specific chemotaxis assays. The Mean Migration Indices to the respective imprinted odor dilutions were determined. Chemo-attractive responses segregate into three stably inherited levels: half crossed worm populations display the maternal behavior (R-), 1/4 lost the negative imprinting and behave as naive (R), while 1/4 show MMI values significantly lower than maternal (R- - < R- < R, **p < 0.01).

Using chemotaxis assays, Mean Migration Indices (MMI), which represent the respective attractive values of a specific odorant for a worm population, were determined for naive and for exposed worm populations. For each odorant, **R** values represents the MMI of naive, **R+** values the enhanced MMI of positively imprinted, while **R-** values the decreased MMI of negatively imprinted worms.

For each odorant, **R+** and **R-** hermaphrodites were crossed with their respective **R+** and **R-** males. From each cross, at least 20 worm populations were obtained from single F1 individuals. F2 and following generations (F2 to Fn) of all populations were submitted to chemotaxis assays.

Crossing positively imprinted **R+ x R+** worms generates three worm populations displaying three different levels of chemo-attractive responses to the respective imprinted odorants (**Figure 6a**). Half progeny behave as the imprinted maternal (**R+**), a quarter behave as naive (**R**), while the remaining quarter acquired chemo-attraction levels **R++** significantly higher than the imprinted maternal behavior **R+**.

The negative imprinting behavior follows a similar segregation pattern in crossed progenies (**Figure 6b**). Crossing negatively imprinted **R- x R-** worms generates three worm populations displaying three different levels of chemo-attraction. Half progeny behave as maternal **R-**, one quarter recover a naive behavior (**R)**, while for the remaining quarter, chemo-attraction was lower than maternal **R-**, so that these worms even exhibit repulsion instead of attraction to the imprinted odors (**R--**).

Worm populations with five different levels of attraction to a single dilution of three different odorant molecules can be thus generated after crossing stably imprinted worms of both sexes. Strikingly enough, once generated, each of the five chemo-attractive behavior, from **R++** to **R--**, is stably transmitted in the self-fertilized hermaphrodites progeny.

A main conclusion can be drawn from **Figure 6** data: the segregation pattern of behavioral changes in the crossed progeny of imprinted worms reproduces a Mendelian segregation pattern expected from crossing stable heterozygotes. As schematically depicted in **Figure 7**, only stable monoallelic alterations can theoretically explain how five distinct levels of chemo-attractive response to a single odorant could be encoded at the genomic level.

**Figure 7:**
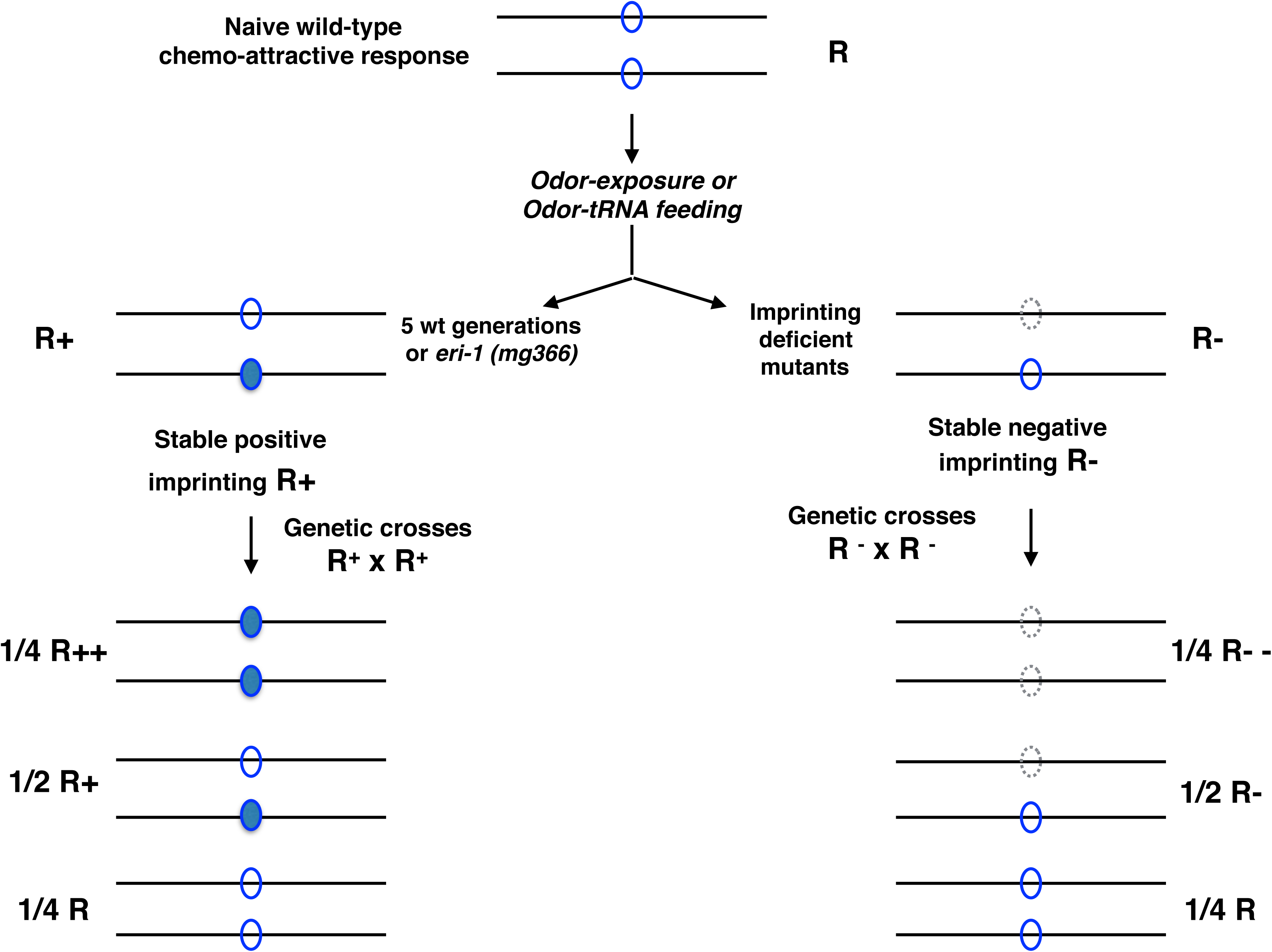
Stable imprints encode stable « behavioral heterozygosity » in the self-fertilized progeny. R represent the chemo-attractive responses of naive worms. Stable positively imprinted R+ were obtained either after five exposed N2 generations or after a single exposed generation of *eri-1 (mg366).* Stable negatively imprinted R- were obtained after a single exposed generation of imprinting deficient *elpc-1 (ng10)* mutants. The observed quantitative (1/2/1) Mendelian segregation of behavioral responses indicates heterozygosity is stably maintained in the self progeny of imprinted hermaphrodites.

Assuming the odor-specific chemo-attractive behavior **R** in naive worms is encoded by a single gene, environment-induced imprinting would modify a single allele of this gene. In the case of positive imprinting, a monoallelic alteration increases chemo-attraction (**R+**), while a different monoallelic alteration inhibits chemo-attraction in negatively imprinted worms (**R-**). In out-crossings, maternal and paternal imprinted and non-imprinted alleles contribute to segregation of the behavioral phenotypes in the progeny. Accordingly, half progeny remains heterozygote, respectively **R+** or **R-**, 1/4 inherit the two parental naive non-imprinted alleles and become homozygotes **R**, while 1/4 inherit the two parental imprinted alleles and become homozygotes, respectively **R++** or **R--**.

Both kinds of imprints would be semi-dominant, as the behavioral phenotypic effects of gain of function positive imprints or of loss of function negative imprints received from both parents are additive.

The initial positive or negative imprinting triggers were single 1/300 dilutions of the respective odorant molecules. The five stable MMI shown in **Figure 7a and 7b** represent worms responses to the imprinted 1/300 dilutions.

In order to evaluate to what extent chemo-attractive responses are modified beyond the imprinting odor dilution, chemotaxis assays were performed using higher and lower dilutions. Panels A and B of **Figure 8** show the responses of respectively naive **R**, positively **R+** and **R++** and negatively **R-** and **R--** CI 1/300 imprinted worms to a 1/100 to 1/900 range of CI dilutions. Panels C and D shows the responses of the **R**, **R+**, **R++**, **R-** and **R--** BA 1/300 imprinted worms to the same range of BA dilutions.

**Figure 8:**
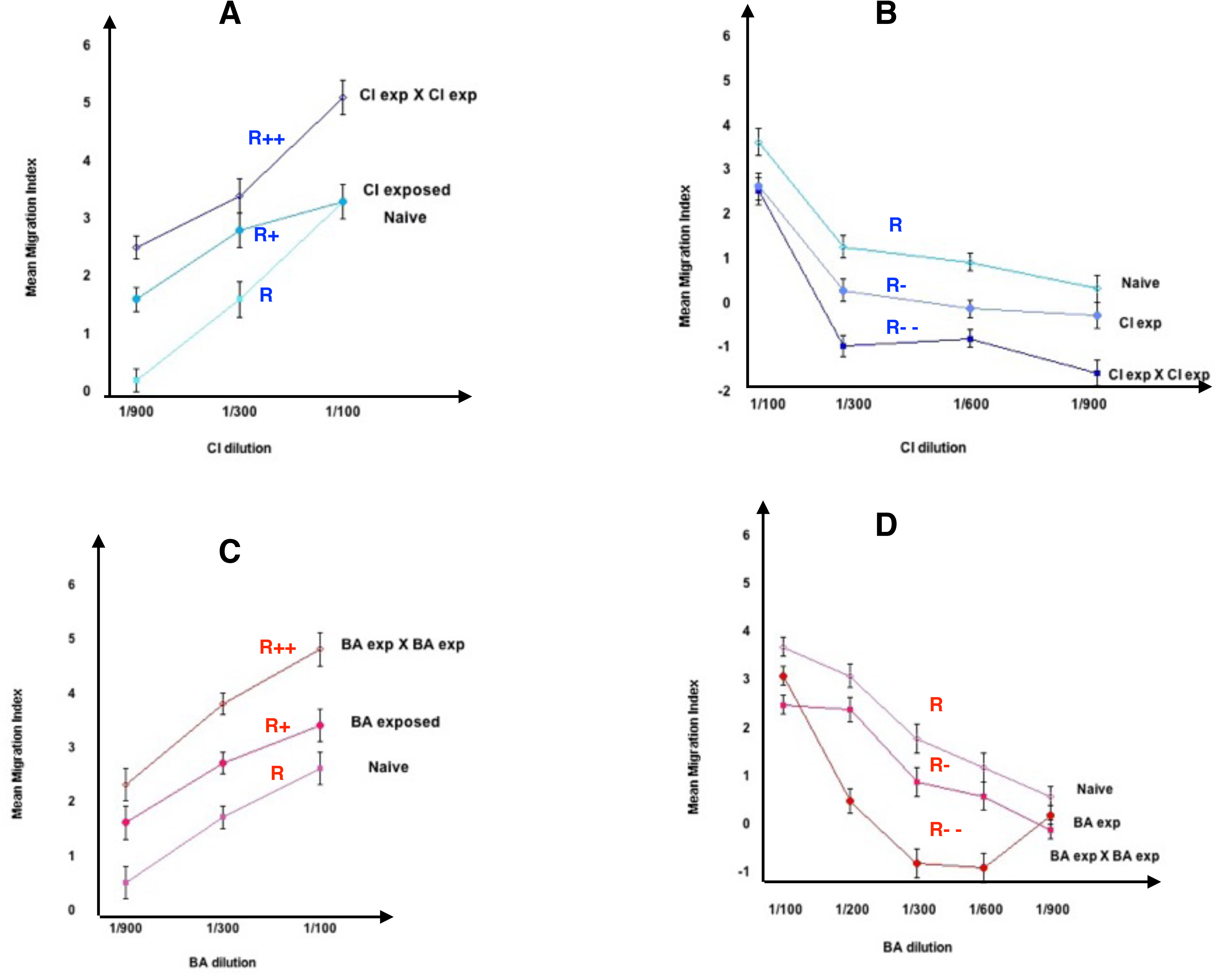
Odor dilution range of modified chemo-attractive responses. A) Chemo-attractive responses (MMI) to the 1/100, 1/300 and 1/900 dilutions of citronellol (CI) of N2 naive (R), of stably CI 1/300 imprinted N2 (CI exposed or R+), and of the crossed progeny stably inherited CI 1/300 imprinted N2 (CI exp X CI exp, R++). B) Chemo-attractive responses (MMI) to the 1/100, 1/300, 1/600 and 1/900 dilutions of citronellol for *elpc-1 (ng10)* naive (R), of CI 1/300 exposed *elpc-1 (ng10)* (CI exp, R-) and of the crossed progeny of stably inherited CI 1/300 imprinted *elpc-1 (ng10)* (CI exp X CI exp, R--). C) Chemo-attractive responses (MMI) to the 1/100, 1/300 and 1/900 dilutions of benzaldehyde (BA) of N2 naive (R), of stably BA 1/300 imprinted N2 (BA exposed, R+), and of the crossed progeny of stably inherited BA 1/300 imprinted N2 (BA exp X BA exp, R++). D) Chemo-attractive responses (MMI) to the 1/100, 1/300, 1/600 and 1/900 dilutions of benzaldehyde (BA) for *elpc-1 (ng10)* naive (R), of BA 1/300 exposed *elpc-1 (ng10)* (BA exp, R-) and of the crossed progeny of stably inherited BA 1/300 imprinted *elpc-1 (ng10)* (BA exp X BA exp, R--).

**Figure 8** dose-response curves indicate that, except for the **R++** worms, positive and negative imprinting respectively enhances or decreases attractive values for lower, but not higher, odor dilutions than the 1/300 imprinting dilution.

The **Figure 7** model postulates the existence of odor-specific genes, each directing chemo-attraction to a limited dilution range of an attractive odorant molecule. The two alleles of a putative « Citronellol attraction» gene and of another putative « Benzaldehyde attraction» gene would be on, respectively, « cici » and « baba » states in naive worms.

Early olfactory experience is adding monoallelic marks, producing the imprinted CIci or BAba states, driving the **R+** enhanced attractive response. Crossing stable CIci or BAba heterozygotes generates a Mendelian ratio (1/4) of CICI or of BABA homozygotes, encoding the **R++** behaviors.

To further confirm « behavioral heterozygosity » and additive effects of stable imprints on worms behavior, crossings between doubly imprinted worms were carried out. Crossed populations were isolated from independent crossing experiments. Results are presented in the form of a classical Punnett square (**Table 2**). The combination of **CI** and BA response levels generated nine different behavioral phenotypes: **R**R, **R**R+, **R**R++, **R+**R, **R+**R+, **R+**R++, **R++**R, **R++**R+, and **R++**R++.

**Table 2:**
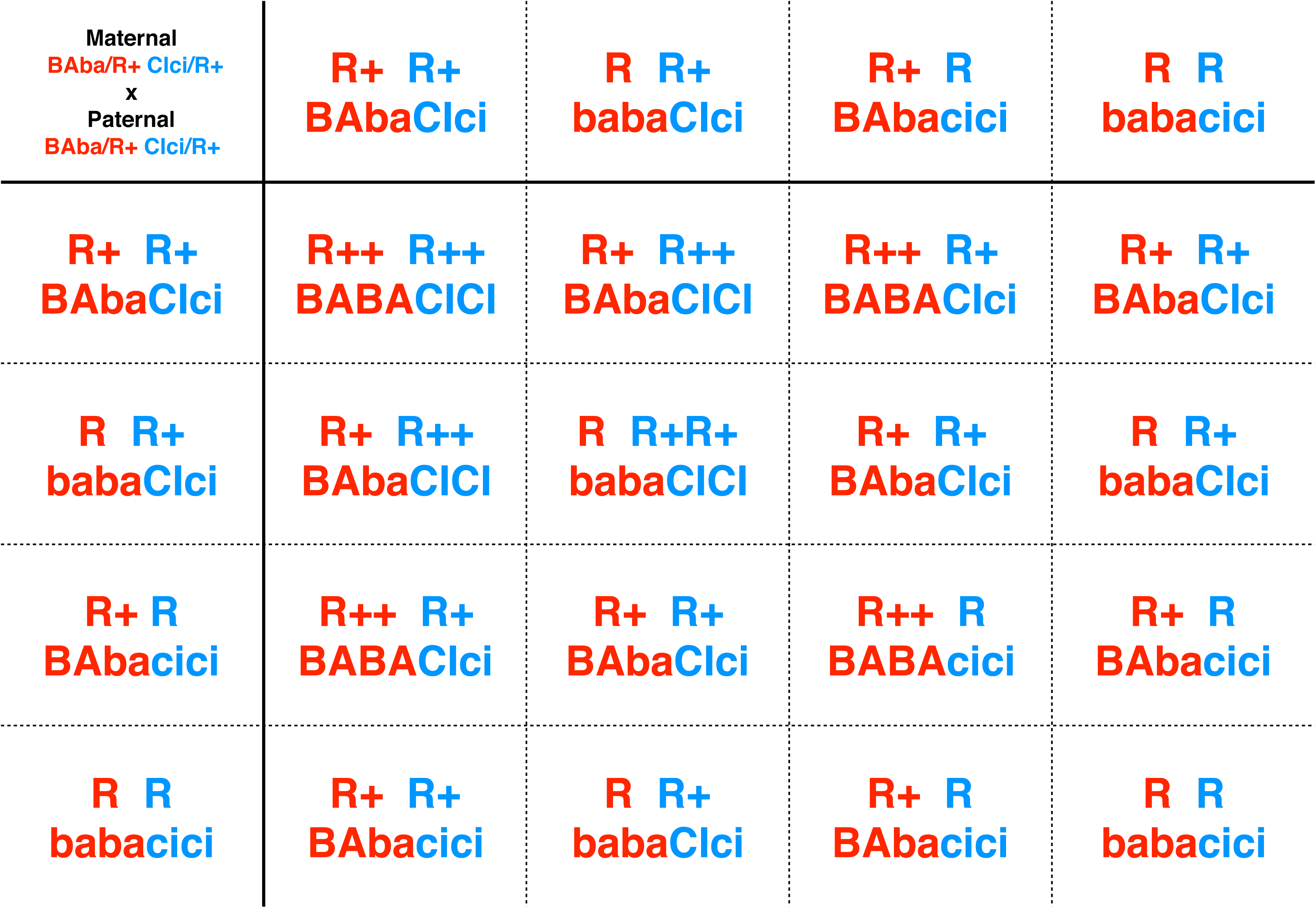
Segregation of CI 300 and BA 300 chemo-attractive values in the progeny of crossed doubly CI + BA imprinted worms (R+ x R+). Punnett square representation of stable behavioral responses (R) segregated from crossing hetero-specifically imprinted worms. The indicated genotypes postulate the existence of a putative gene encoding CI 300 attractive responses and a putative gene encoding BA 300 attractive responses.

| Maternal<br>BAba/R+ Clci/R+<br>x<br>Paternal<br>BAba/R+ Clci/R+ | R+ R+<br>BAbaClci | R R+<br>babaClci | R+ R<br>BAbacici | R R<br>babacici |
| --- | --- | --- | --- | --- |
| R+ R+<br>BAbaClci | R++ R++<br>BABACICI | R+ R++<br>BAbaCICI | R++ R+<br>BABAClci | R+ R+<br>BAbaClci |
| R R+<br>babaClci | R+ R++<br>BAbaCICI | R R+R+<br>babaCICI | R+ R+<br>BAbaClci | R R+<br>babaClci |
| R+ R<br>BAbacici | R++ R+<br>BABAClci | R+ R+<br>BAbaClci | R++ R<br>BABAcici | R+ R<br>BAbacici |
| R R<br>babacici | R+ R+<br>BAbacici | R R+<br>babaClci | R+ R<br>BAbacici | R R<br>babacici |

The nine combinatorial behavioral phenotypes might be respectively encoded by the nine combinations of putative genotypes - respectively babacici, babaCici, babaCICI, BAbacici, BAbaCIci, BA-baCICI, BABAcici, BABACIci and BABACICI - predicted from crossing BabaCIci x BAbaCIci double heterozygotes.

The observed frequency of each combination follows the Mendelian rules. For each individual odorant, attractive values segregate, as predicted from crossing heterozygotes, as 1/4 naive, 1/2 maternal, and 1/4 maternal added to paternal values. One quarter of the worms inherit the maternal double imprint **R+**R+ (BAbaCIci), while 1/16th of the worm populations behave as naive **R**R (babacici) « homozygotes » and another 1/16th as **R++**R++ (BABACICI) « homozygotes ». A mean of 1/8th of worms display either a **R**R+, a **R+**R, a **R+**R++ or a **R++**R+ behavior.

Segregation of behavioral phenotypes in the cross progeny of doubly imprinted is thus as expected from crossing double heterozygous « genotypes ». As already indicated in **Figure 7**, all behavioral or « allelic » combinations described in **Table 2** are stably transmitted over generations in the self progeny.

All together, the data obtained using genetic approaches, suggests olfactory experiences acquired from odor-exposure or odor-tRNA feeding, leave an imprint on a single allele of, presumably, a single member of the olfactory receptor (OR) gene family (***Robertson and Thomas, 2006***).

The molecular identity of the gain-of-function positive and the loss of function negative imprints are unknown, but, as opposed to mutations in DNA sequence, both would establish and maintain stable heterozygosity in hermaphrodite animals. The behavior of homozygous worms generated through crossing suggest imprints have quantitative additive effects on the assignment of attractive values to chemo-attractive odorant molecules. The five different levels of attraction toward a single odorant (**Figure 6 and 8**) might therefore result from five different expression levels of the same OR gene.

## DISCUSSION

Using worm feeding on biochemically purified molecules, the regulatory role of Alanine tRNAs molecules on the *C. elegans* chemo-attractive behavior has been previously demonstrated (***Fernandes de Abreu, 2020***). Different Ala tRNAs isomers may control both the implementation of innate and the environment-modulated chemo-attractive responses.

The co-culture and food conditioning experiments carried out in the present study show that the same behavioral regulations are observed without artificial addition of purified Ala-tRNAs to worm food. These evidences lead to the conclusion that regulatory Ala-tRNAs might be naturally released and present in worms environment.

Feeding bacterial food conditioned by the presence of naive worms restore chemo-attraction in the *elpc-3* mutants. With the exception of *elpc-3* mutants, all tested worms, including the chemo-attraction and imprinting mutants, may release the *elpc-3* rescuing developmental Ala-tRNA, suggesting its constitutive secretion.

Odor-specific forms of Ala-tRNA (odor-tRNAs) are made in response to early larval odor-stimuli. Odor-tRNAs can be considered as molecular mnemonics of worms early olfactory experiences. They increase adult chemo-attractive responses via feeding. Worms carrying mutations that inactivate the transduction of olfactory messages, or had lost odor-responsiveness, do not release odor- tRNAs.

Increased responses due to imprinting can be horizontally commmunicated from odor-exposed to naive commensals - sharing the same food -, suggesting odor-tRNAs are secreted and spread in the vicinity of odor-exposed animals.

Worm-to-worm communication of imprinting needs the intestinal dsRNA specific SID-2 molecule. SID-2 is the entry point of environmental dsRNA, required for the systemic spreading of RNA interference gene silencing (***Winston et al., 2007***; ***McEwan et al., 2012***). This argues in favor of the presence of regulatory tRNAs in worms environment. Imprinting and its inheritance indeed require a number of genes belonging to the RNAi pathways.

In *C. elegans,* exogenous RNAi, triggered by the external addition of dsRNA to worm food, is widely used experimentally to efficiently silence the target genes with complementary sequences (***Timmons and Fire, 1998***).

In plants and insects, the exo-RNAi machinery has been linked to antiviral immunity, as it is able to target and process dsRNA viral genomes (***Ding, 2010***). In *C. elegans*, a role for exogenous RNAi as an potential viral defense mechanism has been only revealed after the discovery of dsRNA viruses infecting nematodes in the wild (***Félix et al., 2011; Ashe et al., 2013***). The way RNAi acts as a antiviral defence in *C. elegans* might not however fully overlap with the canonical gene silencing RNAi triggered by the addition of gene complementary dsRNA in worms environment. Viral dsRNA genomes do not display sequence homology with the worm genome, suggesting RNAi triggered by non-self sequences might differ from RNAi triggered by self sequences. Amongst noticeable differences between the two mechanisms, antiviral defence seems not systemic nor transgenerationally transmitted (***Ashe et al., 2015***).

Besides immunity, exogenous RNAi triggered by gene-complementary dsRNAs might thus support additional biological functions in the current life of *C. elegans* nematodes. Previous studies demonstrated that RNAi indeed mediates behaviors in responses to environmental signals (***Juang et al., 2013; Sims et al., 2016; Posner et al., 2019***). However, in these examples, the initial dsRNA triggers and the targeted genes remain unknown.

Odor-tRNAs are molecular memories and vectors of early olfactory experiences. As inferred from the data presented here, they are indeed able to horizontally transfer the imprinting behavior to naive. However, spreading memories in order to share passed experiences with neighbouring naive might not be the actual biological function of odor-tRNAs secretion in natural conditions.

All larvae sharing the same olfactory environment would be imprinted by the same olfactory cues, suggesting there is no need for worm-to-worm horizontal transmission. Moreover, it seems hard to assign a communication function to the released developmental form of Ala-tRNA. Every worm indeed develope a chemo-attractive behavior without the need of developmental Ala-tRNA emitters in the vicinity.

What could be therefore the biological function supported by the external release of regulatory Ala tRNAs and what could be the mechanisms by which this single molecule, via feeding RNAi, could regulate *C. elegans* chemo-attractive responses?

The hypothesis discussed below seems more consistent with the evidence presented here and with the currently available knowledge.

Interaction of odorant molecules with specific olfactory receptors (OR) expressed by the chemosensory neurons is, up to now, the only admitted molecular basis for odor-specific responses. Externalized regulatory Ala-tRNAs, instead of inter-individual communications, may be part, through Ala-tRNA processing by the RNAi silencing machinery, of a feed-back loop mechanism by which a worm epigenetically modulates the expression of its own OR genes.

This hypothesis raises however a number of challenging questions:

### 1) Regulatory Ala-tRNAs would be part of a putative odor-sensitive RNA secretome

Exosomal RNAs, as yet undescribed in *C. elegans*, is thought to mediate different forms of communications, including intercellular, interindividual and cross-species transfer of information. In parasitic nematode species, extracellular vesicles (ECV) containing RNA secretomes, mostly made of small non-coding RNAs, including highly aboundant tRNA fragments, may play a role in crossspecies parasite to host communications (***Buck et al., 2014***; ***Quintana et al., 2019***).

In *C. elegans*, ECV are released by all ciliated sensory neurons, that include the chemo-sensory neurones mediating chemo-attraction. It has been suggested that these ECV, whose molecular composition is yet unknown, could function in worm to worm communications (***Wang et al., 2014***). Ala-tRNAs might possibly belong to a RNA secretome released by odor-responsive chemo-sensory neurons ECV.

### 2) Regulatory Ala-tRNAs may be processed into siRNAs

Imprinting and its inheritance involves a number of RNAi genes: the systemic RNAi SID-2, needed for RNA uptake from the environment, members of the Dicer complexe that process dsRNAs into primary 21-nt long small interfering RNAs (exo-siRNAs), the RdRP RRF-3 and the ERI-1 RNase, both involved in the biosynthesis of secondary 26-nt endo-siRNAs, refered as 26G as they mostly start with a guanine nucleotide (***Han et al., 2009***; ***Billi et al., 2014***).

As suggested by the behavioral phenotype of worms carrying the *rrf-3 (pk1426)* or the *eri-1 (mg366)* mutations (**Figure 3**), 26G secondary siRNAs seem to play an essential role in regulating the imprinting behavior. The *eri-1 (mg366)* worms are 26G RNAs depleted, and the RNA dependent RNA polymerase (RdRP) activity of RRF-3 is necessary for 26G RNA accumulation.

By contrast, the two *rrf-1 (pk1417)* and *rrf-1 (ok589)* mutants are wild-type for imprinting and imprinting inheritance, suggesting the RdRP activity of RRF-1, generating 22G secondary siRNAs, is not involved (***Billi, 2014***).

The *rrf-3 (pk1426)* worms do not imprint, neither positively nor negatively (**Figure 3, Table 1**), while without a paternal ERI-1 activity, imprinting escape from erasing and become stably fixed in the progeny.

Interestingly, 26G RNAs are distributed into two gender-specific populations, according to which Argonaute protein they bind: ALG-3 and ALG-4 in the spermatogenic gonad, ERGO-1 in the oogenic gonad (***Gent et al., 2010***).

Ala-tRNAs processing by the Dicer complexe would generate primary siRNAs made of Ala-tRNA homologous fragments (21-nt si-tRNAs). These putative sitRNAs could further serve as template for the synthesis of secondary putative 26G si-tRNAs by RRF-3.

The *C. elegans* genome encodes 19 different Argonautes proteins, each of them binding distinct classes of small RNAs with some specificity. RNA providing the target specificity, Argonaute- RNAs compexes are guided to complementary sequences. To which Argonaute binds Ala tRNA derived siRNAs remains to be discovered.

### 3) Ala-tRNA RNAi may epigenetically control the expression of olfactory receptor genes

The permanent behavioral alterations due to positive or negative imprinting may result from odor-induced, RNAi dependent, up or down epigenetic regulations of odor-specific olfactory receptor (OR) monoallelic gene expression.

In *C. elegans*, about 1500 predicted genes encode a putative superfamily of OR which belongs to the G protein-coupled receptors class (***Troemel et al., 1995; Robertson and Thomas, 2006***). How worms control OR genes expression in chemo-sensory neurons is still unknown.

The regulation of OR genes expression seem to be based on the same general principles in mammalian species and in Drosophila (***Iyengar et al., 2010; Magklara, 2011; Lyons, 2013; Ferreira, 2014; Golovin, 2019; Jafari, 2021***). They can be summarized as follows. Olfactory experience du-ring critical periods of early development influence the implementation of future chemo-sensory responses. Expression of OR genes, or of single OR gene alleles in mammals, would depend of receptor activation by odorants present in the early environment. The functional differenciation of chemo-sensory neurones is terminated by OR expression, which is epigenetically controlled by heterochromatin switches, initiating transcription. Feedback mechanisms will further inhibit the expression of all other OR genes in the same cell.

Mammalian and insect OR gene promoters contain repressive heterochromatin H3K9me2 histone marks. OR genes repression or expression seem at least, to involve differential methylation of H3K9 into H3K9me2 or H3K9me3, depending on the activity of species-specific methylases and demethylases.

RNAi is a conserved mechanism of gene regulation which had been linked to chromatin modifications *(**reviewed by*** ***Seroussi, 2021***). Strinkingly, it has been proven that RNAi support epigenetic inheritance by coupling siRNAs to H3K9 methylation states in different species (***Klosin, 2017***; ***Yu, 2018).*** Environmentally-triggered changes in chromatin states therefore provide a general mechanism for epigenetic transmission of information between generations (***Burton, 2011; Duempelmann, 2020***).

It could be inferred from the data presented here, that different chemo-attractive values might be based on differential odor-specific OR expression levels in worm chemo-sensory neurones. Stable positive and negative imprints would result from odor-tRNA RNAi induced, stably inherited, addition of monoallelic repressive or permissive epigenetic marks.

It has to be noted however that only positive monoallelic imprinting would actually regulate worms behavior in natural conditions. Negative imprinting only occurs in animals genetically impaired for positive imprinting. Homozygote imprinted states might be thus rare in the wild, due to the scarcity of sexual reproduction in *C. elegans*. Non-naturally occuring phenotypes can be generated through experimental designs, reflecting a broad extension of phenotypic plasticity, beyond ecologically or physiologically relevance, as already discussed (***Baugh, 2020***).

### 4) Ala-tRNA siRNAs might target the extra-TFIIIC sites present on the C. elegans genome

Ala-tRNA derived siRNAs could only target tRNA complementary sequences, that is tDNA genes and transcription independent extra-TFIIIC sites.

tDNA genes are transcribed into tRNAs by a complex associating Polymerase III and the two TFIIIC and TFIIIB transcription factors. TFIIIC binds to two conserved internal promoters called the A and B boxes, present in all tRNA sequences. TFIIIC bound to TFIIIB form the pre-initiation complex recognized by Polymerase III.

Extra-TFIIIC (ETC) sites are chromosomal locations independent of tDNA genes, carrying the TFIIIC interacting A and B boxes.

tDNA genes and ETC sites play a role in the organisation of chromatin domains as insulators. In the Yeast, they can moreover silence the neighboring Polymerase II transcribed genes (***Noma, 2006; Donze, 2012***).

The current knowledge strongly pleads in favor of ETC sites instead of tDNA genes as the potential targets of Ala tRNA siRNAs to control OR genes expression (***Stutzman, 2020)***:

- Most of the 504 *C. elegans* ETC sites are found on the autosomes (chromosomes I to V in the worm), and not on the X sexual chromosome, as opposed to the 525 tDNA genes (respectively 486 versus 18 on the X and 323 versus 202 on the X).
- The worm ETC sites are over-represented on chromosome V (40% of the total) while OR genes are strongly clustered and also over-represented on chromosome V, while quite absent on the X chromosome (***Robertson, 2006***).
- The worm ETC sites are clustered on repeats and intergenic loci on the distal arms of autosomes, located close to, but not overlapped with, H3K9me2 and H3K9me3-enriched regions, the epigenetic marks involved in epigenetic OR gene regulation in other species.
- By contrast to tDNA genes, worm ETC sites are almost all (88%) located within LEM-2 subdomains present at the nuclear periphery, which regulate large scale chromosomal spatial organisation. As in other species, TFIIIC binding to ETC sites on worm genome may demarcate the different active and repressive chromatin states. A similar distribution for ETC sites and OR genes on worm genome would be compatible with a regulatory function of ETC sites on OR genes expression via dynamic chromatin structural changes.

RNAi processing of constitutive and environment-induced Ala tRNAs into small ETC sites binding RNAs would support behavioral diversification through remodelling the spatial organization of odor responsive genes.

## Material and Methods

### Worm strains

Otherwise indicated, all worm strains used in this study (N2 Bristol (wild-type), N2 carrying the syls179 (lin-4::YFP) transcriptional fusion, *odr-1 (n1936), tax-2 (p671), tax-4 (p678), rrf-3 (pk1426), rrf-1 (ok589)*, *rrf-1 (pk1417)*, *eri-1 (mg366), eri-1 (mg366) carrying a myo-3::gfp transgene; sid-1 (pk3321), sid-1 (qt2), rde-1 (ne300), rde-1 (ne219), drh-2 (ok951), rde-2 (ne221), mut-7 (pk204), sid-1 (qt2), elpc-3 (ok2452),* and *daf-22 (m130)),* were provided by the Caenorhabditis Genetics Center (CGC). The *drh-1 (tm1329)* and *elpc-3 (tm3120)* mutant strains were obtained from Dr S. Mitani (National Bioresource Project, Tokyo, Japan). The *sid-2 (qt13)* mutant was from Dr C. Hunter. The elongator mutant *elpc-1 (ng10)* was from Dr J. A. Solinger, while the *rde-4 (ne337)* and *rde-4 (ne299)* mutants, isolated by the Dr C. Mello laboratory, were obtained from Dr O. Hobert.

### Odor-exposure

Benzaldehyde, β-citronellol or Isoamyl alcohol (Sigma-Aldrich) were diluted as described in water. Odor-exposures were done by suspending a 4 µl drop of these dilutions on the lids of worm culture dishes at least during 24 hours from the egg stage at 20°C, covering the critical plasticity period corresponding to the first 12 hours of post-hatch development (***Remy and Hobert, 2005***).

### Chemo-attraction assays

Chemotaxis assays were fully described previously in ***Fernandes De Abreu (2020)***. Mean migration Indices (MMI) were compared using the student t test for unpaired data with unequal variance of the KaleidaGraph program. The Mean Imprinting Indices (MII) represent the difference between Mean Migration Indices of two different populations for a specific odorant dilution.

### Food conditioning

NGM agar culture plates were loaded with a single drop of OP50 liquid culture using a 1 ml pipette tip (Sarstedt). After drying the surface of E. coli spots were approximately 3 cm^2^ .

### Stability of CI-tRNA on OP50 spots

In order to obtain highly purified Alanine tRNA (UGC) molecules from CI 1/300 exposed N2 worms, we used the Streptavidin Sepharose^TM^ High Performance beads (GE Healthcare, 17-5113-01) coupled to the 3’-biotinylated tDNA Ala (TGC)-3’ specific probe (5’-TGGAGG-TATGGGGAATCGAACCCCAGACTTCTCGCAT-3’). The purification procedure of CI-tRNAs has been fully described in **Fernandes De Abreu (2020).**

## Acknowledgments

We thank Craig Hunter for the *sid-2 (qt13)* mutant and Craig Mello for the *rde-4 (ne337)* and *rde-4 (ne299)* mutants. The elongator mutants *elpc-1 (ng10)* and *elpc-3 (tm3120)* were obtained thanks to Jachen A. Solinger via Grazia Malabarba. Thanks to the National Bioresource Project, Tokyo, Japan, which is part of the International C. elegans Gene Knockout Consortium, for the *elpc-3 (tm3120)*, and the *drh-1 (tm1329)* strains. Other strains used in this study were provided by the Caenorhabditis Genetics Center funded by the National Institutes of Health (NIH) Office of Research Infrastructure Programs (P40 OD010440). We thank Wormbase for providing valuable databases.

Thanks to Valérie Grandjean for stimulating discussions and incentive suggestions. This work was supported by the “Agence Nationale de la Recherche” ANR-12-Bioadapt-0022 grant.

