## Supplemental Figure and supplemental Table for "An environment to genome control loop using RNA interference processing of secreted tRNAs may regulates the *C. elegans* chemo-sensory behavior"

**Supplemental Figure Legend**

**The *elpc-3* rescuing developmental Ala tRNA is constitutively released**

Blue bars
Naive N2 were used as Emitters, while the Elongator sub-unit 3 *elpc-3 (tm 3120)* and *elpc-3 (ok 2452)* mutants were used as Receivers, as described in Figure 4. Worms carrying the *elpc-3 (tm 3120)* and *elpc-3 (ok 2452)* mutations were fed either on unconditioned food or on Naive N2 conditioned food (Naive N2 Emitters, *elpc-3* mutants Receivers).
Mean Migration Index to CI 1/300 of *elpc-3* Receivers was determined as described.

Red bars
Naive Elongator sub-unit 3 *elpc-3 (tm 3120)* and *elpc-3 (ok 2452)* mutants were used as Emitters, while N2 worms were used as Receivers, as described in Figure 4. Naive N2 were fed either on unconditioned food or on Naive *elpc-3* conditioned food (Naive *elpc-3* Emitters, N2 Receivers). MMI to CI 1/300 of N2 Receivers was determined as described.

**Supplemental Table: Stability of biochemically purified CI-tRNA on OP50 bacterial lawns.**

| **Purified<br>Cl-tRNA (UGC)** | **No delay** | **24 hours delay** | **48 hours delay** | **72 hours delay** | **96 hours delay** |
|---|---|---|---|---|---|
| **1 ng** | **Imprinted** | **Imprinted** | **Imprinted** | **Imprinted** | Naive |
| **100 pg** | **Imprinted** | **Imprinted** | **Imprinted** | Naive | Naive |
| **10 pg** | **Imprinted** | **Imprinted** | Naive | Naive | Naive |
| **1 pg** | **Imprinted** | **Imprinted** | Naive | Naive | Naive |
| **100 fg** | **Imprinted** | **Imprinted** | Naive | Naive | Naive |
| **10 fg** | **Imprinted** | **Imprinted** | Naive | Naive | Naive |
| **1 fg** | **Imprinted** | Naive | Naive | Naive | Naive |

**Table 1: Stability of Cl-tRNA $^{Ala}$ (UGC) on bacterial lawns.**

**Supplemental Figure**

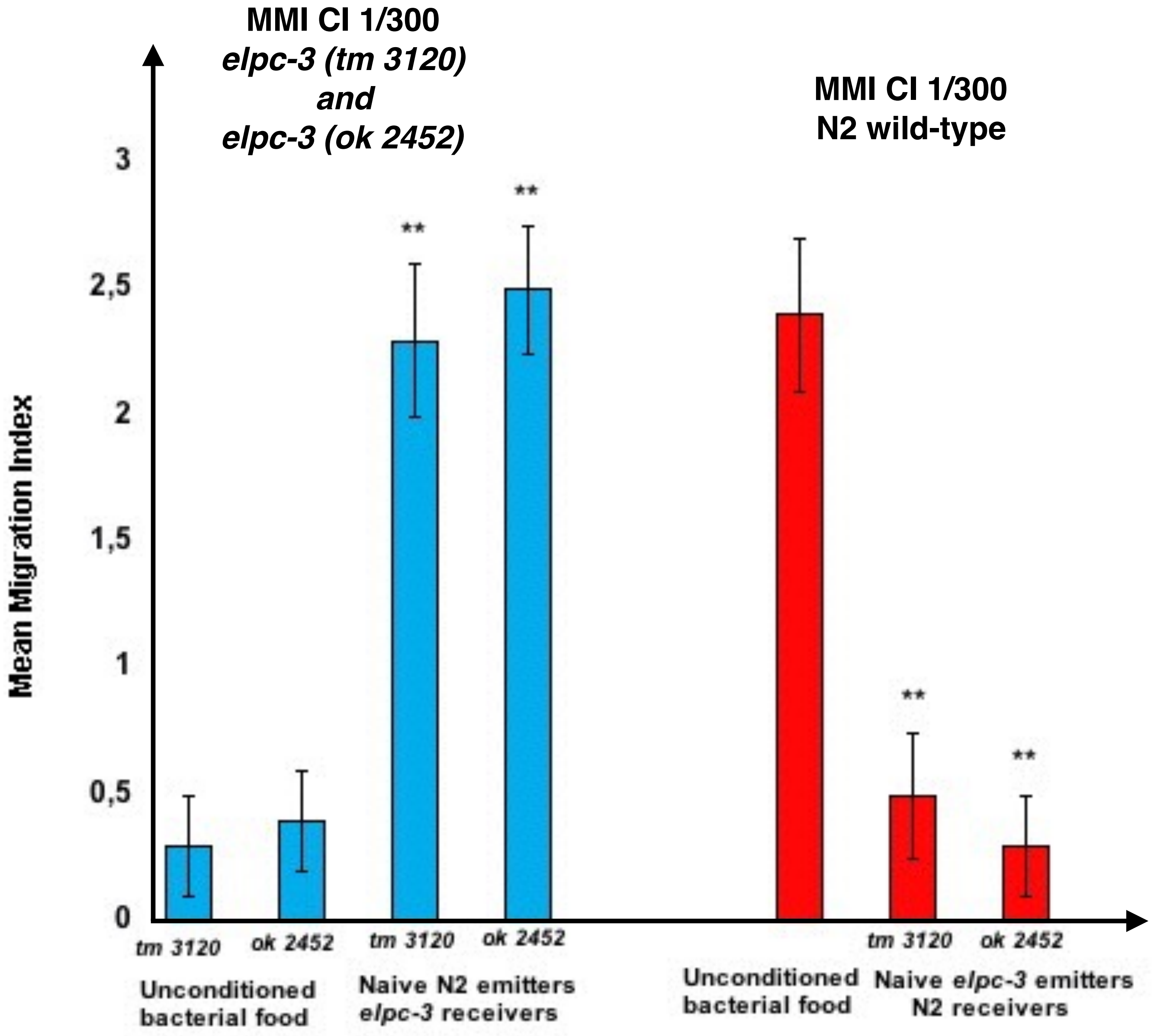
